# Structural identification of the RY12 domain of RyR1 as an ADP sensor and the target of the malignant hyperthermia therapeutic dantrolene

**DOI:** 10.1101/2024.10.21.619409

**Authors:** Kookjoo Kim, Huan Li, Qi Yuan, Zephan Melville, Ran Zalk, Amédée des Georges, Joachim Frank, Wayne A. Hendrickson, Andrew R. Marks, Oliver B. Clarke

## Abstract

Malignant hyperthermia (MH) is a life-threatening pharmacogenetic condition triggered by volatile anesthetics, which activate pathogenic RyR1 mutants. The small molecule therapeutic dantrolene has long been used to treat MH. However, the binding site and mechanism of dantrolene remain unclear. Here, we present cryo-EM structures of RyR1 bound to dantrolene and the MH trigger agent 4-chloro-m-cresol (4CmC), revealing the dantrolene and 4CmC binding sites in atomic detail. Dantrolene binds stacked with ATP or ADP in the RY12 domain at the corner of the receptor, inducing a conformational change in this domain which is allosterically coupled to pore closure. Functional analyses revealed that ATP or ADP was required for dantrolene inhibition, and a single point mutation that disrupts the peripheral ATP binding site abolished ATP/ADP-dependent dantrolene inhibition. Strikingly, in the absence of dantrolene, this site selectively binds two ADP molecules, suggesting a possible role in ATP/ADP ratio sensing.

## Introduction

RyR1 is an intracellular Ca^2+^ channel that plays a key role in excitation-contraction coupling (EC coupling) by mediating efflux of Ca^2+^ from the SR to the cytoplasm, triggering skeletal muscle contraction^1,2^. Gating of RyR1 is regulated by many ligands, ions and binding partners, for example, metal ions such as Ca^2+^ and Mg^2+^; endogenous and exogenous ligands such as adenosine triphosphate (ATP), adenosine diphosphate (ADP), caffeine^3^; and proteins, such as calstabin1 (Cs1, also known as FKBP12)^4,5^ and calmodulin (CaM)^6,7^. The C-terminal domain (CTD) of RyR1 contains binding sites for Ca^2+^, ATP/ADP and caffeine/xanthine^8^. In addition to the central ligand binding hub, the RY12 domain located at the corner of the RyR1 tetramer has been recently reported to bind a class of RyR1/RyR2 targeting therapeutics known as “Rycals”, in conjunction with ATP.^9^

Stringent regulation of intracellular Ca^2+^ release is essential, since uncontrolled Ca^2+^ release can have severe pathophysiological consequences, as exemplified by malignant hyperthermia (MH). Malignant hyperthermia susceptibility (MHS) is a pharmacogenetic condition most commonly linked to RyR1 mutations, where patients experience muscle rigidity and a rapid rise in body temperature when exposed to certain volatile anesthetics (MH trigger agents). These compounds, which do not affect wild-type RyR1 at therapeutic doses, abnormally activate mutant RyR1, leading to a hypermetabolic state, and the increased rate of compensatory ATP hydrolysis leads to hyperthermia, rhabdomyolysis, acidosis, and often death if untreated.

Dantrolene, the only approved therapeutic agent for the treatment of MH, inhibits RyR1-mediated Ca^2+^ release^10–12^, but the precise binding site of dantrolene and its molecular mechanism of action have remained elusive, although experiments with tritiated azidodantolene showed that it lies somewhere within the first 1400 amino acids of the receptor^12^, and that ATP, Mg^2+^ and CaM may potentiate the inhibitory effect of dantrolene on RyR1^13–16^.

The binding sites and mechanism of action of MH trigger agents (including halothane anesthetics and 4-chloro-m-cresol) also largely remain to be elucidated. Perhaps the best mechanistically studied MH trigger agent mechanistically is 4-chloro-m-cresol (4CmC), which is not an anesthetic but was originally used as a preservative agent in succinylcholine^17^, prior to being identified as a potent RyR1 activator. 4CmC also increases Ca^2+^ affinity at the high-affinity Ca^2+^ binding site in the core of the receptor^18^. Two key residues for activation of RyR1 by 4CmC, Q4020 and K4021, have been identified^19^, but in structures of RyR1 K4021 projects from the surface of the core solenoid into the solvent, while Q4020 lines a mostly hydrophobic cavity in the core of the domain, with no obvious entry points, leaving the exact binding site of 4CmC and mechanism unresolved.

In this study, we have obtained multiple high-resolution single particle reconstructions of the RyR1-Cs2-CaM complex in the presence of various ligands, including dantrolene and 4-CmC, in order to probe the binding sites and potential mechanism of MH triggers, MH therapeutics, and adenine nucleotides. We have localized the binding site of dantrolene to the RY12 domain at the corner of the receptor, where it binds as a ternary complex with ADP or ATP, similar but distinct from the binding mode of Rycal therapeutics^9^, accompanied by closure of the cleft between the two subdomains, and remodeling of the interprotomer interface. The high local resolution of 2.5Å in this region after local processing has allowed us to confidently assign atomic details of the interaction. In single-channel lipid bilayer recordings, ATP or ADP is required for RyR1 inhibition by dantrolene, and a single point mutation (W882A) which disrupts the ATP binding site abolishes ATP/ADP-dependent dantrolene inhibition. In the absence of dantrolene, we show that the RY12 domain selectively binds two ADP molecules, even in the presence of a substantial excess of ATP, and adopts a conformation that is strikingly similar to the dantrolene-bound conformation.

In structures with 4CmC present, we have confirmed that 4CmC interacts directly with Q4020, occupying a cryptic pocket in the inaccessible center of the core solenoid domain. This site is separated from the high affinity Ca^2+^-binding site by a single layer of alpha-helices, providing a potential explanation for how 4CmC can modulate RyR1 sensitivity to Ca^2+^. We also identified three auxiliary 4CmC binding sites, in the transmembrane (TM) domain, the EF-hand domain, and the RY12 domain.

## Results

We initially prepared cryoEM grids of purified RyR1-Cs2-apoCaM complex in the presence of a combination of three RyR1 agonists: ATP, caffeine, and 4CmC to maximize open probability so that we could investigate the effect of dantrolene binding to RyR1 in both the open and closed states. Mg^2+^ was included due to evidence that it may potentiate dantrolene binding to RyR1^16^. Four conditions were investigated: Dantrolene/ATP/4CmC/caffeine/Mg^2+^ (DanATP), ATP/4CmC/caffeine (ATP4CmC), Dantrolene/ADP/4CmC/caffeine/Mg^2+^ (DanADP), and ADP/4CmC/caffeine/Mg^2+^ (ADP4CmC) to recapitulate both the open and closed states of RyR1 in the presence of the three activators including 4CmC, an MH trigger, with and without the presence of dantrolene in the buffer.

CryoEM data collection and processing statistics are summarized in Table 1 Detailed information on purification conditions and concentrations of the ligands used in each condition is summarized in Methods, and tabulated in Table 2. Domain boundaries of rabbit RyR1 are summarized in Table 3. A representative micrograph, 2D class averages, and 3D reconstruction are shown in Figure S6. Data processing workflows are described in Methods and Figures S7 & S8. The peripheral cytosolic domains of RyR1, including RY12, are highly mobile, and as such they remain poorly resolved in consensus reconstructions using a global mask for refinement, necessitating symmetry expansion and careful local refinement and classification in order to improve the interpretability of density in the peripheral regions of the receptor.

**Table 1.**
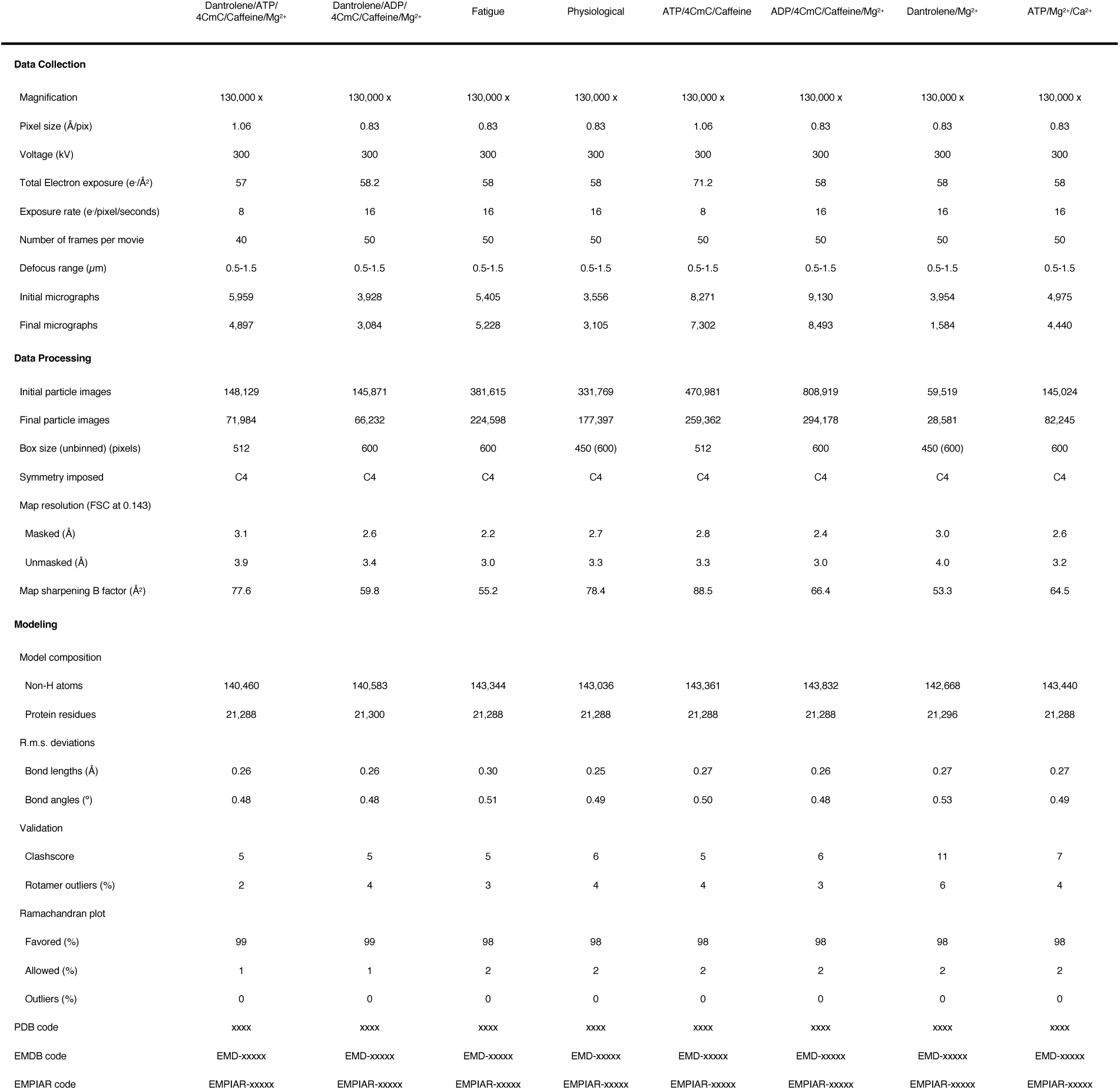
CryoEM data collection, refinement and validation statistics.

**Table 2:** Conditions and ligand concentrations for cryoEM grid preparations.

| Name | Dantrolene | ATP | ADP | 4CmC | Caffeine | Free Mg<sup>2+</sup> | Free Ca<sup>2+</sup> |
| --- | --- | --- | --- | --- | --- | --- | --- |
| ATP4CmC | - | 2 mM | - | 2 mM | 5 mM | - | 100 nM |
| DanATP | 0.1 mM | 2 mM | - | 2 mM | 5 mM | 0.2 mM | 100 nM |
| ADP4CmC | - | - | 0.2 mM | 2 mM | 5 mM | 0.2 mM | 100 nM |
| DanADP | 0.1 mM | - | 0.2 mM | 2 mM | 5 mM | 0.2 mM | 100 nM |
| Fatigue | - | 1.2 mM | 0.2 mM | - | - | 1 mM | 100 nM |
| Physiological | - | 7 mM | 0.01 mM | - | - | 1 mM | 100 nM |
| DanMg | 0.1 mM | - | - | - | - | 0.2 mM | 100 nM |
| ATPMgCa | - | 2 mM | - | - | - | 1 mM | 30 μM |

**Table 3:** Description of the different local masks used for focused local refinement.

| Mask Name | Domains | Residue Span (rabbit RyR1) |
| --- | --- | --- |
| Corner | SPRY1-RY12-SPRY2-SPRY3 | 628-1656 |
| Core-Pore | CSol-TM-Pore-CTD | 3667-5037 (for all 4 chains) |
| JSol | Cs2, JSol, BSol (partial), CaM, SCLP | 1657-2365, 3624-3682 + Cs2 & CaM |
| BSol | BSol-RY3&4 | 2366-3623 |
| NSol | NTD-NSol | 1-627 |

### Dantrolene binds to the RY12 domain in an ATP/ADP dependent manner

After masked local refinements with a local mask around the three SP1a/ryanodine receptor (SPRY) domains (SPRY1, SPRY2, and SPRY3) and RY12 which includes amino acid residues 628-1656 in rabbit RyR1, dantrolene was identified in the local EM maps with either ADP (Figure 1) or ATP (Figure S1) binding in a stacking manner between the two α-helices in the RY12 “clamp” at 2.3 Å and 2.6 Å local resolution, respectively. In this cleft, dantrolene and ADP/ATP are flanked by W996, which stacks with the hydantoin moiety of dantrolene, and W882, which stacks with the adenine moiety of ADP/ATP. The positively charged amino acid residues R1000, R1020, R886, R897, and H904, line the binding pocket to accommodate terminal phosphates of the adenine nucleotides (Figure 1B&C). Dantrolene binds to the inner side of the cleft, while ADP is positioned on the outer side of the left, in contrast to Rycals such as ARM-210, which have been shown to bind on the outer side of the cleft, stacking with an inner ATP molecule positioned where dantrolene is located here. We then investigated whether the presence of ATP or ADP is required for RyR1 inhibition by dantrolene. We incorporated the purified RyR1-Cs2-CaM complex into proteoliposomes for planar lipid bilayer single-channel recordings (see Methods for details). 100µM dantrolene decreased the open probability (P_o_) of RyR1s that were first activated by 1 µM Ca^2+^ and either 2 mM ATP or ADP, but in contrast dantrolene addition did not decrease P_o_ in the absence of ATP or ADP (Figure 1D). The current-time traces of single-channel recordings are shown in Figure S2. Based on the RyR1-dantrolene-ATP/ADP ternary complex structures and the observation that ATP or ADP was required for dantrolene inhibition, we hypothesized that the binding of ATP or ADP at the dantrolene binding site (as opposed to the central ATP/ADP binding site in the CTD) is required for the binding of dantrolene and the inhibitory effect of dantrolene. ATP and ADP both stack directly with the indole ring of W882, leading us to suppose that mutation of this residue would weaken or abolish ATP/ADP binding at the peripheral site. Thus, RyR1-W882A was tested by single-channel bilayer recording. RyR1 with W882A substitution was still activated by ATP and Ca^2+,^ but no longer showed ATP/ADP-dependent inhibition by dantrolene (Figure S2). Thus, dantrolene inhibition requires cooperative binding of either a molecule of ATP or ADP bound at W882 in the RY12 cleft in order to inhibit RyR1. These results are consistent with structural observations, which show no evidence of dantrolene binding in the absence of ATP/ADP (Figure S3).

**Figure 1.**
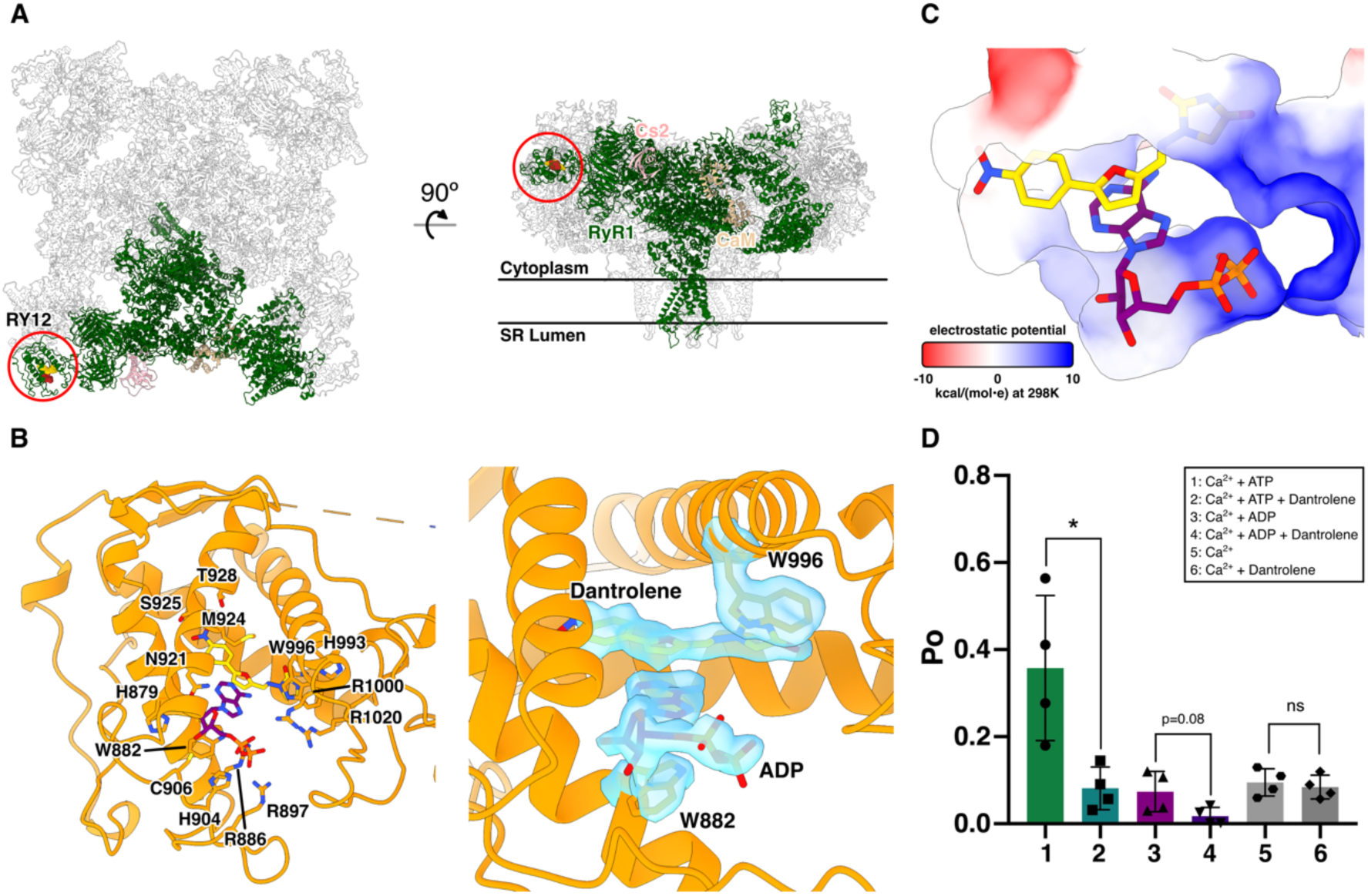
Dantrolene binds in the RY12 domain (RY12) with ADP or ATP. **A.** The location of the RY12 domain in an RyR1 protomer (green). Dantrolene (gold) and ADP (maroon) are shown in the side and top view ribbon representations of the RyR1-Calstabin2 (FKBP12.6; pink)-Calmodulin (tan) complex. **B.** Left: Dantrolene, ADP, and the side chains in the RY12 domain. Right: Model fit for W996, dantrolene, ADP, and W882 in a density-modified cryo-EM map which is refined and reconstructed from a sub-particle extraction of RyR1 particles re-centered on the RY12 domain. **C.** Dantrolene and ADP in the surface representation of the RY12 binding site colored by Coulombic electrostatic potential. **D.** Quantification of single channel lipid bilayer recordings of purified RyR1 in response to 100 µM dantrolene. 2 mM ATP or ADP was first added before dantrolene. The *cis* [Ca^2+^], representing the cytoplasmic compartment, was kept at 1 *µ*M for all conditions (n=4, data point presented as mean ± SD). *p=0.018 by Student’s T-test.

### The RY12 domain selectively binds ADP in the absence of dantrolene

In the case of the ADP4CmC (ADP/4CmC/caffeine/Mg^2+^) dataset, which lacked dantrolene, two molecules of ADP were found to bind to RY12 in a stacked manner bridged by Mg^2+^, accompanied by closure of the RY12 cleft similar to that observed in the presence of dantrolene, whereas the ATP4CmC (ATP/4CmC/caffeine) dataset did not show RY12 cleft closure or evidence of ordered nucleotides binding in the RY12 domain. This prompted us to investigate whether RY12 could selectively bind ADP in the presence of an excess of ATP, at physiological concentrations that are present in a myocyte. We solved structures in two conditions of different [ATP] and [ADP]: a physiological ATP/ADP condition and a muscle fatigue condition. During muscle fatigue, [ATP] is decreased because of increased rate of ATP hydrolysis, and ADP is generated, increasing [ADP] in myocytes. Reconstructions of RyR1 in the muscle fatigue state yielded a 2.2 Å map from a consensus refinement before masked local refinement. In the local RY12 map from the fatigue dataset, a stack of two molecules of ADP was identified in RY12, with no bound ATP evident, despite the 6:1 ATP:ADP ratio. The interaction between the phosphate groups of the two stacked ADP molecules is mediated by a Mg^2+^ ion (Figure 2). By contrast, under conditions mimicking the resting state, with a much larger excess of ATP, we no longer see evidence for ADP binding at high occupancy in the RY12 domain, although a small sub-class can still be isolated where ADP is bound. The fact that the ADP-bound state of RY12 closely resembles the dantrolene-bound state of RY12 strongly suggests that ADP may be inhibiting RyR1 by binding at this site, and raises the intriguing possibility that the RY12 domain may act as a sensor for the ATP:ADP ratio in muscle, acting to dampen RyR1 activity when cellular ATP levels are depleted.

**Figure 2.**
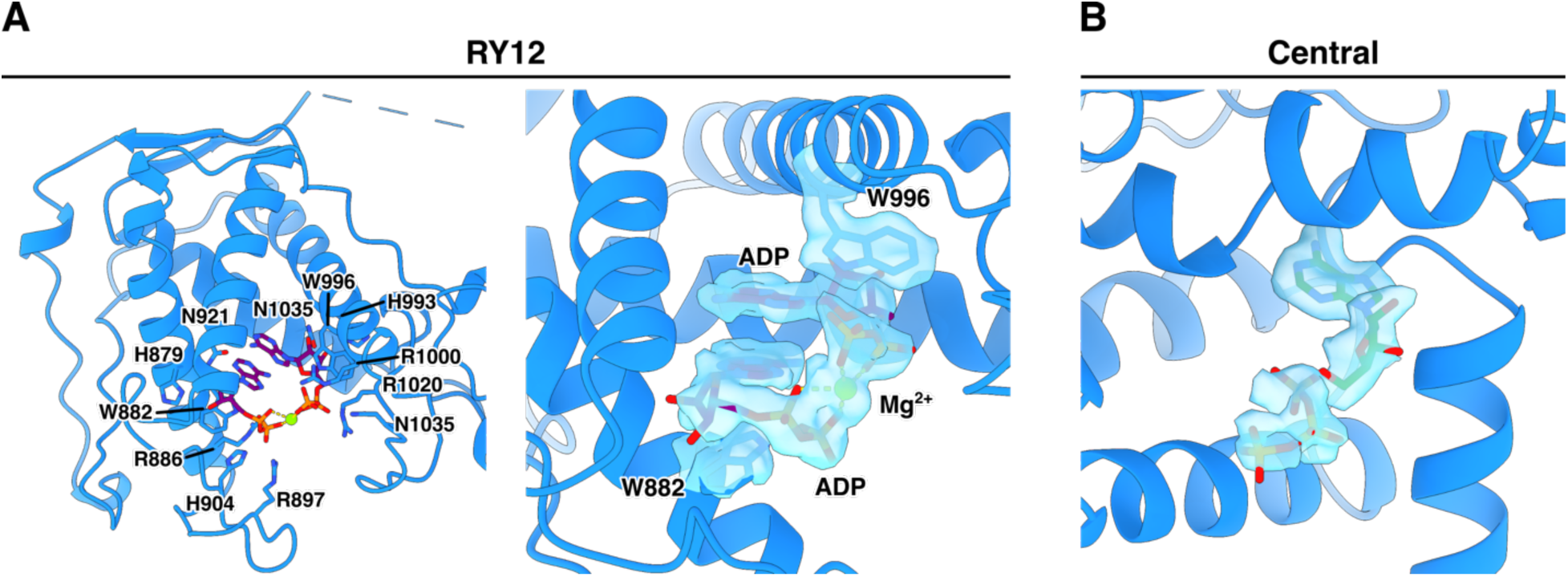
Two molecules of ADP stack in RY12 with Mg^2+^ in [ATP] = 6 x[ADP] in buffer. **A.** Coulomb potential density extracted from the cryoEM map of the RY12 binding site of the RyR1 complex in buffer representing the [ATP] and [ADP] in muscle fatigue. **B.** Coulomb potential density extracted from the fatigue RyR1 map of the C-terminal Domain (CTD) binding site for ATP.

### 4CmC binds to Q4020 in a cryptic pocket within the core solenoid

In all datasets of RyR1 complexed with 4CmC (Dan/ATP, Dan/ADP, ATP/4CmC, & ADP/4CmC), three 4CmC binding sites per RyR1 monomer were identified: One primary site located within the center of the core solenoid (CSol) domain, interacting with Q4020, which has previously been shown to be essential for 4CmC activation, and two auxiliary sites – one located 11Å away, on the surface of CSol at the base of the EF-hand domain, and a second auxiliary site within the membrane embedded part of the receptor, binding to the outside of the transmembrane domain (Figure 3). In the two adjacent binding sites within CSol and the EF-hand domain, Q4020 and Q4133 directly interact with the two 4CmC molecules (Figure 3B & S5), and multiple hydrophobic amino acid residues line these sites (Figure 3C&D). The primary 4CmC molecule bound to Q4020 is buried in a cryptic pocket without a direct access pathway from the solvent, implying that conformational change of CSol is required for ligand binding. The transmembrane 4CmC is wedged in between the S5 and the P helices of one RyR1 protomer and pore lining S6 helix of the adjacent RyR1 protomer (Figure 3D). The two 4CmC binding sites in CSol are in proximity to the pore-forming domain and the C-terminal domain (CTD), where the ATP, Ca^2+^, and caffeine/xanthine bind. Additionally, in the local RY12 map of ATP/4CmC dataset, a molecular model of 4CmC could be fit in the map with an ATP molecule (Figure S4), identifying a third auxiliary site. This, again, is comparable to the binding modes of Dan/ADP and Dan/ATP, in which the ATP molecule stacking on top of W882 sidechain is conserved, and the other member of the molecular stack (4CmC in this case) is seemingly interchangeable with an exogenous ligand such as dantrolene, although the fact that 4CmC is not observed to bind here in the presence of dantrolene or ADP suggests that the affinity of 4CmC at this site is likely lower than the other ligands known to bind to RY12.

**Figure 3.**
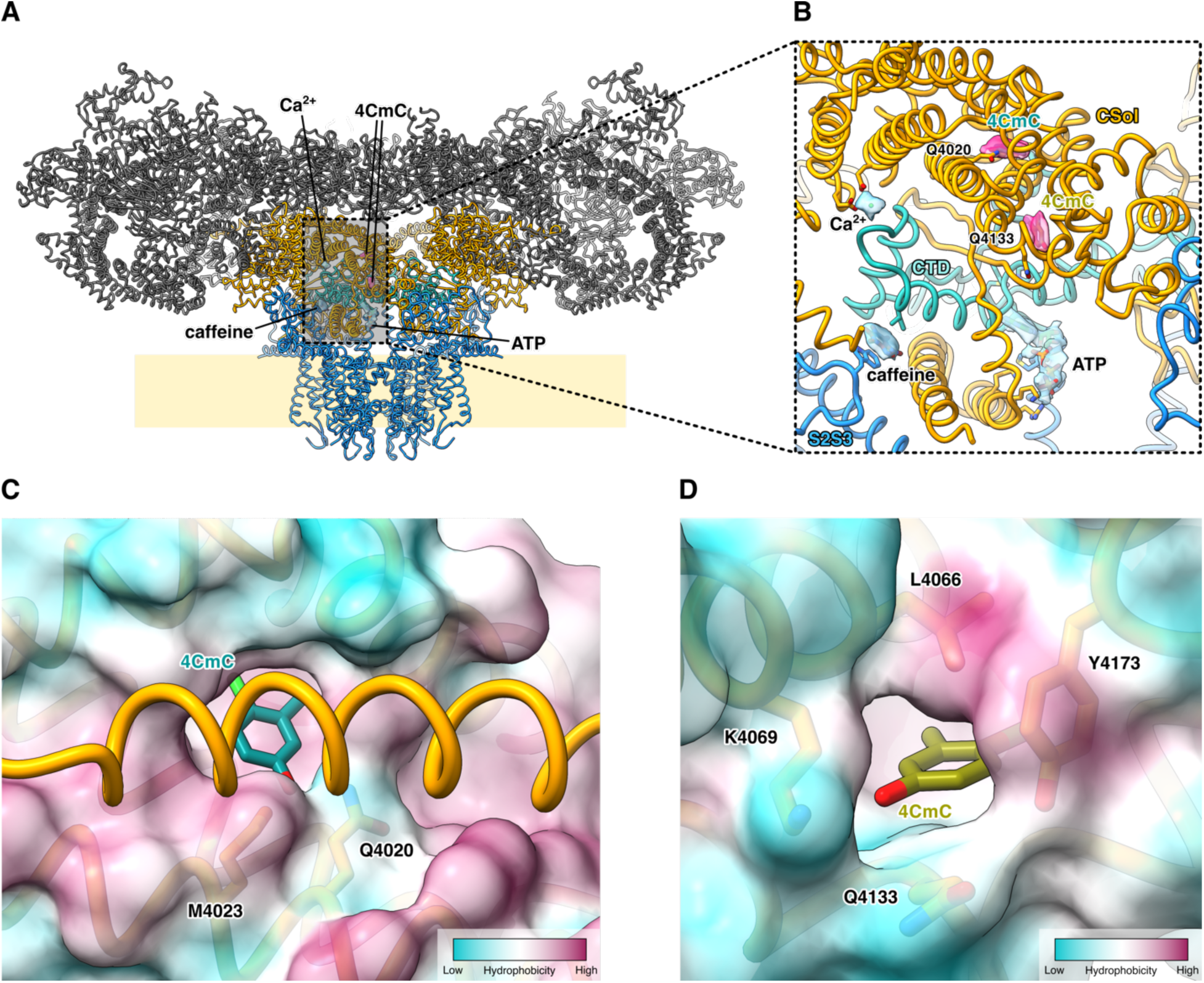
The 4CmC binding sites in the CSol domain of RyR1. **A.** The two 4CmC (magenta) binding sites in CSol (gold) are located adjacent to the CTD (aquamarine) binding sites for ATP, Ca^2+^, and caffeine (sky blue). **C&D.** The surface representation of the RyR1-4CmC model colored by hydrophobicity. The Q4020 site, which is not directly accessible from the buffer, is in a more hydrophobic environment than the Q4133 site is.

### 3D variability analysis reveals altered conformational coupling between BSol and CSol in the presence of dantrolene

After obtaining high-resolution reconstructions from global and local refinements for each dataset, we set out to explore and identify conformational heterogeneity present in the datasets, especially around the core-pore region and the corner region of RyR1. Previously, it has been reported that a downward movement of the BSol domain towards the membrane is coupled to opening of the transmembrane pore of RyR1 and RyR2,^8,20^. In each of the two datasets with dantrolene (Dan/ATP and Dan/ADP), 3D Variability Analysis (3DVA) in *cryoSPARC*^21^ identified a principal component that represents an “inverted” mode of RyR1 conformational coupling, whereby a slight downward movement of the BSol domain is coupled to closure of the transmembrane pore (Figure 4). The Cɑ- Cɑ distances of the hydrophobic gate residue, I4937, were measured as proxies for pore diameters of open and closed states from the normal and the inverted modes. The pore diameters did not significantly differ between the putative open conformations of the normal mode and the inverted mode (Figure 4B).

**Figure 4:**
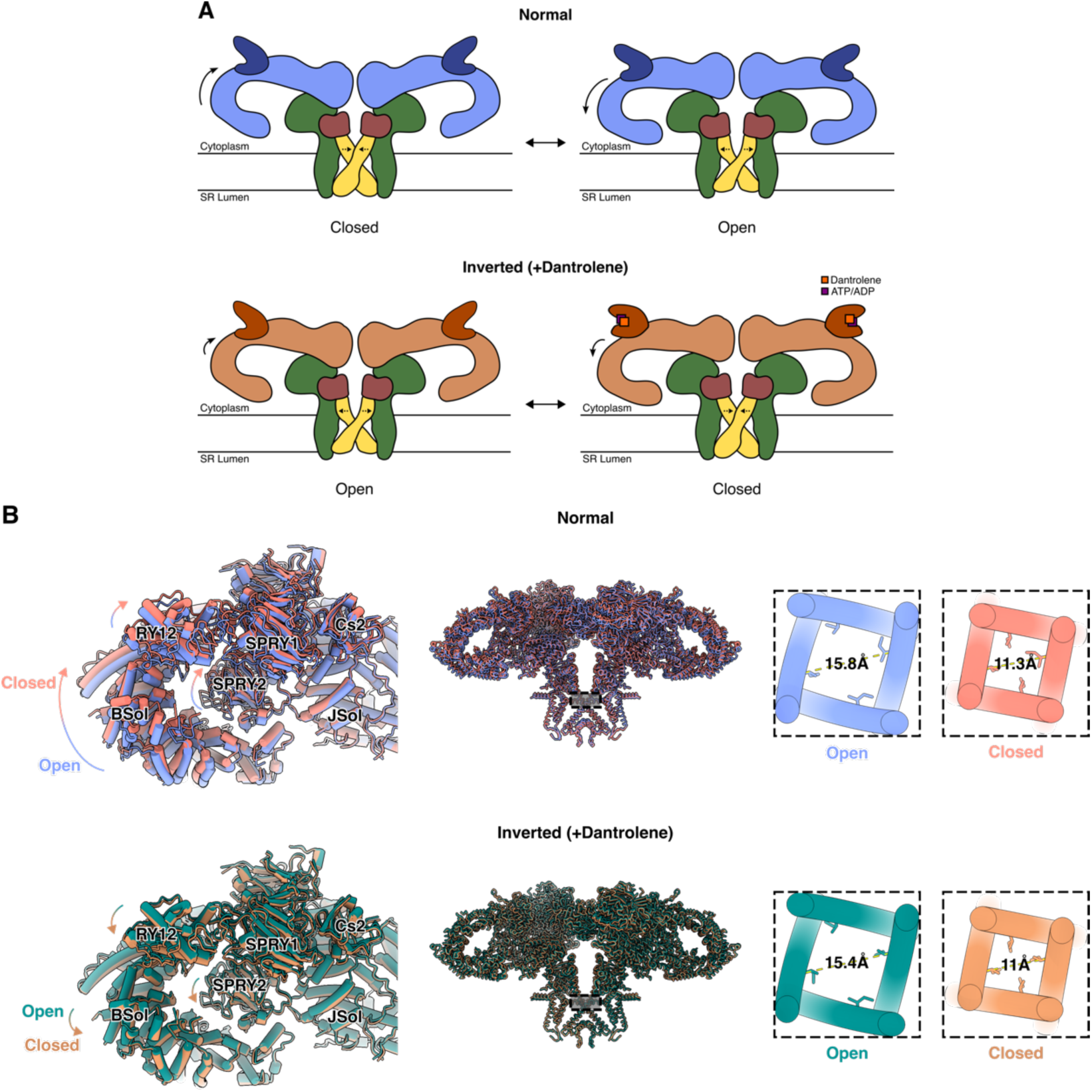
Inverted coupling of BSol position and pore aperture is observed in the presence of dantrolene. **A.** Schematic representations of Normal and Inverted (+Dantrolene) coupling modes. **B.** Neighboring BSol and SPRY domains exhibit an inverted relative domain movement when dantrolene & ATP/ADP are bound in RY12, and the pore diameters (Cɑ distances of I4937 in the pore) are comparable to that of open and closed channels.

## Discussion

### RY12 hosts binding sites for both endogenous ligands and therapeutics

The structural and functional investigation of purified rabbit RyR1 bound to dantrolene, as well as the MH trigger molecule 4CmC, that is presented in this report complement each other and help elucidate the molecular mechanism of dantrolene inhibition of RyR1. With high-resolution (2.6 Å & 2.3 Å) local reconstructions around RY12, we could unambiguously assign dantrolene and ATP/ADP in the local cryoEM maps. Moreover, dantrolene did not inhibit RyR1 without ATP or ADP in planar lipid bilayer recordings (Figure 1D & Figure S2), nor did it bind to RY12 in the absence of adenosine nucleotides (Figure S3) This is consistent with prior photolabeling studies using [^3^H]azidodantrolene to label RyR1 expressed in CHO cells, where the [^3^H]azidodantrolene photolabeling of RyR1 was greatly enhanced in the presence of AMP-PCP, a non-hydrolyzable ATP analog^22^. In terms of the local conformational changes of RY12 upon binding of ligands, when dantrolene and ATP/ADP are bound, the RY12 cleft closes like a clamp to form the binding pocket. It is not fully understood how this local conformational change at the corner of the RyR1 tetramer is propagated to the channel pore, but the proximity of this site to the interprotomer interface suggests that one possibility may be that closure of the RY12 clamp helps stabilize the interprotomer interface, which is often destabilized by MH mutations^23^. Moreover, RyR1s form a checkerboard-like array *in situ* and *in vitro*,^24^ and it has been recently reported that this paracrystalline arrangement is preserved in mouse extensor digitorum longus muscle *in situ* probed by cryo-electron tomography (cryo-ET)^25^. The arrangement of the arrays indicates that RY12 is likely to be involved in interactions between individual channels in the array on the SR membrane, raising the possibility that ligands that stabilize the RY12 domain may also affect coupling between adjacent channels in an array^26^. The observation that in the absence of dantrolene, the RY12 domain seems to selectively bind ADP in an inhibitory conformation, even in the presence of a physiologically relevant excess of ATP, raises the intriguing possibility that the RY12 domain may be acting as an endogenous sensor of the ATP:ADP ratio, dampening RyR1 activity when cellular ATP is depleted, for example during muscle fatigue, thus preventing the cell from entering the hypermetabolic state which can lead to rhabdomyolysis.

### 4CmC binds to a cryptic pocket in the CSol, as well as several auxiliary sites

4CmC is a potent RyR1 activator, initially identified as an MH trigger when it was used as a preservative in the preparation of succinylcholine^17^. Previously, Fessenden *et al*. showed that Q4020 and K4021 are required for RyR1 activation by 4CmC, but the structural details of this interaction were unclear, even after the initial structures of RyR1 were solved^19,27^. Indeed, we observed in the cryo-EM maps that Q4020 directly interacts with one of the two 4CmC molecules in CSol. The primary 4CmC binding site is not readily accessible to the solvent (Figure 3C), and a substantial conformational rearrangement of the cytosolic domain of RyR1 would be required for 4CmC binding. The role, if any of the auxiliary 4CmC binding sites in 4CmC activation of RyR1 remains unclear; However, previous studies on isolated RyR1 using single channel bilayer recordings indicate that 4CmC produces an additive effect on RyR1 activity when it is added simultaneously to both the cytoplasmic (*cis* side of the bilayer) and the luminal (*trans* side of the bilayer) sides, which would be unexpected if the only activating site were located in the cytosol^19^. The identification of a 4CmC binding site in the luminal leaflet of the bilayer may provide a possible explanation for luminal activation by 4CmC, but further studies will be required to show this definitively.

Previous studies have shown that the RY12 is also the binding site for the putative therapeutics known as Rycals^9,28^ which are currently undergoing clinical trials for cardiac and muscle disorders caused by leaky RyR channels^29^. Rycals bind to the RY12 site, also in a ternary complex with a molecule of ATP, and via an allosteric mechanism stabilize the closed state of the channel thereby preventing a pathological leak of intracellular Ca^2+^. In contrast to Dantrolene, Rycals do not inhibit the channel, and stabilize a wider configuration of the RY12 cleft. The fact that both dantrolene and Rycals such as ARM- 210 bind to the same site in an ATP-dependent manner, but on opposite sides of the RY- 12 cleft, suggests a possible complementarity between the two classes of therapeutics, and an explanation for their differing effects upon the channel.

**Figure S1:**
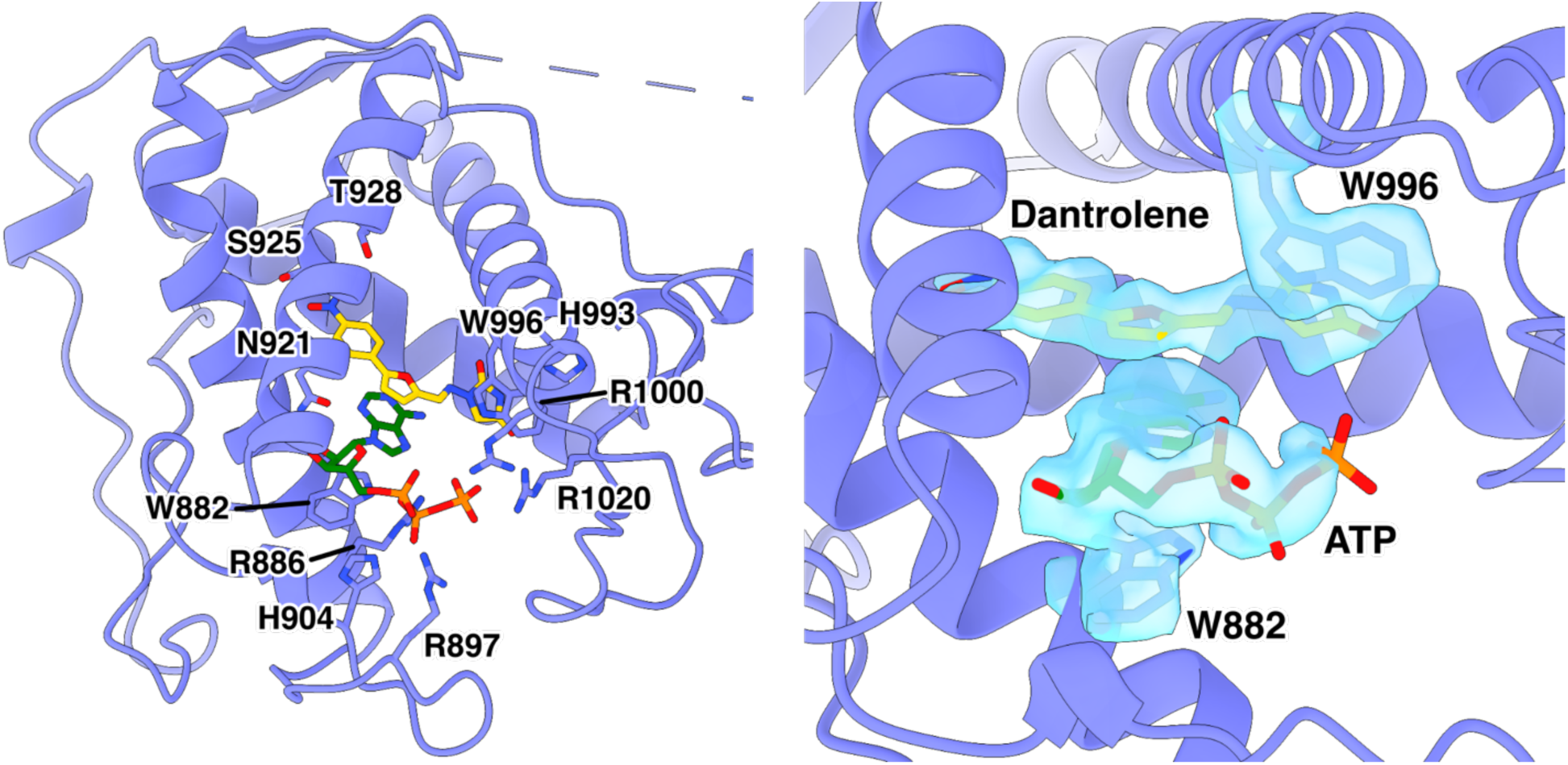
ATP binds in RY12 with dantrolene. The atomic model of RY12 is colored in light purple. Like Dan/ADP, dantrolene and ATP are flanked by W996 and W882.

**Figure S2:**
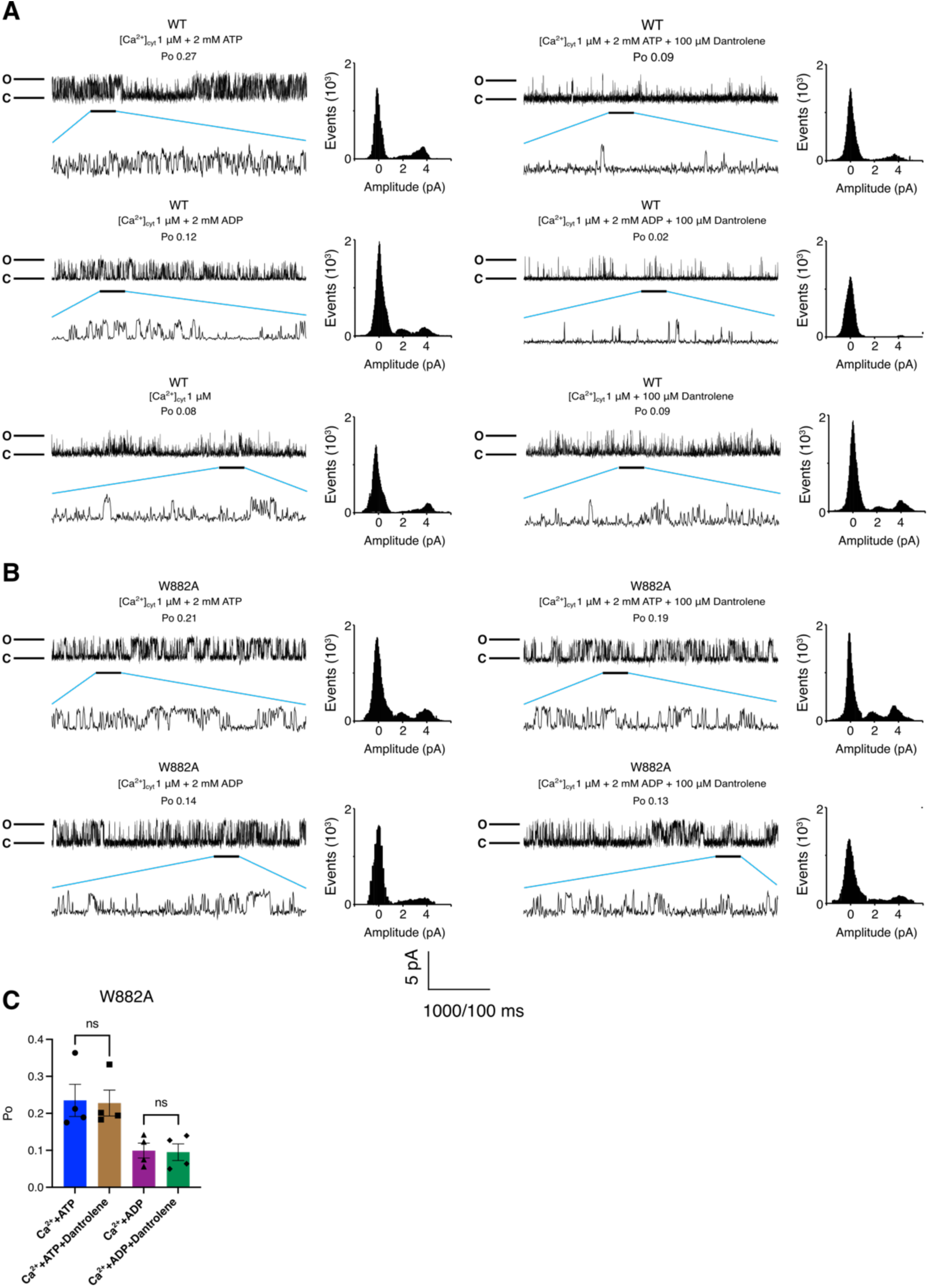
The current traces of wild-type & W882A mutant RyR1 **A.** Single-channel traces of wild-type (WT) RyR1 in a planar lipid bilayer after the addition of 1 *µ*M Ca^2+^ with 2 mM ATP, 1 *µ*M Ca^2+^ with 2 mM ADP, and 1 *µ*M Ca^2+^ alone. Amplitude histograms for each condition are shown by the current traces. The agonist effect of each ligand is abolished by the addition of 100 *µ*M dantrolene. **B.** Single-channel traces of W882A mutant RyR1 in a planar lipid bilayer after the addition of 1 *µ*M Ca^2+^ with 2 mM ATP and 1 *µ*M Ca^2+^ with 2 mM ADP. Amplitude histograms for each condition are shown by the current traces. The agonist effects of ATP and ADP on W882A RyR1 are not abolished by the addition of 100 *µ*M dantrolene. **C.** Quantification of single channel lipid bilayer recordings on W882A mutant RyR1 microsome in response to 100 *µ*M dantrolene. 2 mM ATP or ADP was first added before addition of dantrolene (n=4, data point presented as mean ± SD). The *cis* [Ca^2+^], representing the cytoplasmic compartment, was kept at 1 *µ*M for all conditions.

**Figure S3:**
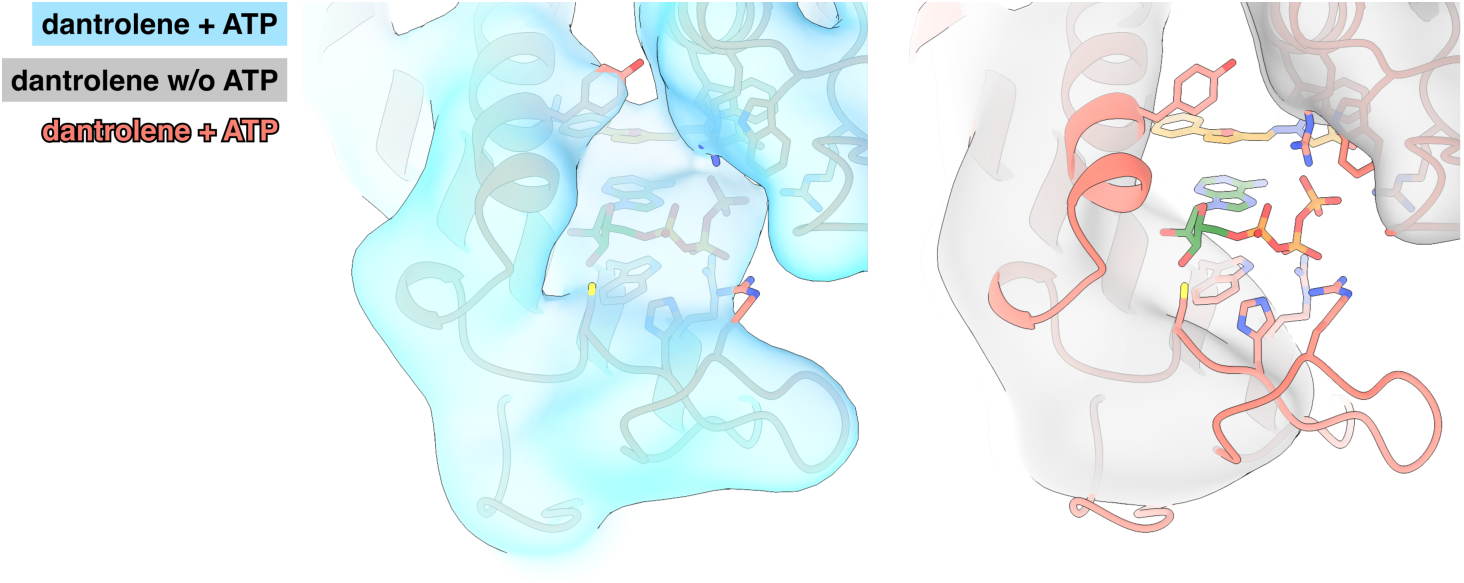
Without ATP, Dantrolene does not bind in RY12 (Dantrolene/Mg^2+^). Lowpass-filtered cryoEM maps from Dan/ATP/4CmC/Caffeine/Mg^2+^ (skyblue) and Dan/Mg^2+^ (grey) datasets. The RY12 domain structure complexed with dantrolene, and ATP is colored in salmon.

**Figure S4:**
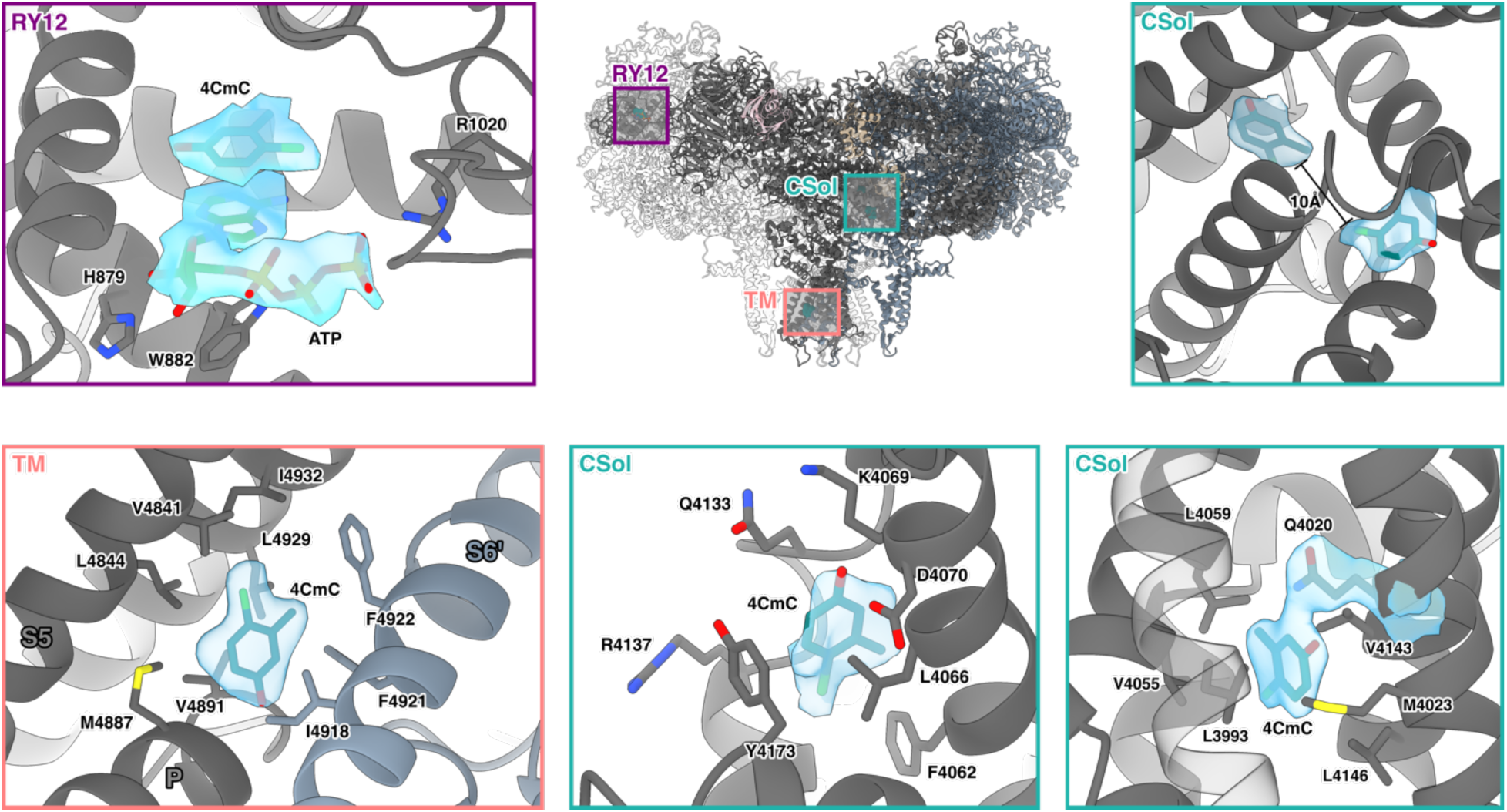
Different 4CmC binding sites on RyR1. 4CmC and ATP bind together in RY12 (purple box). The two 4CmC binding sites in CSol are ∼10Å apart from each other (teal box). 4CmC also can dock in between P and S6 helices in TM (salmon box).

**Figure S5:**
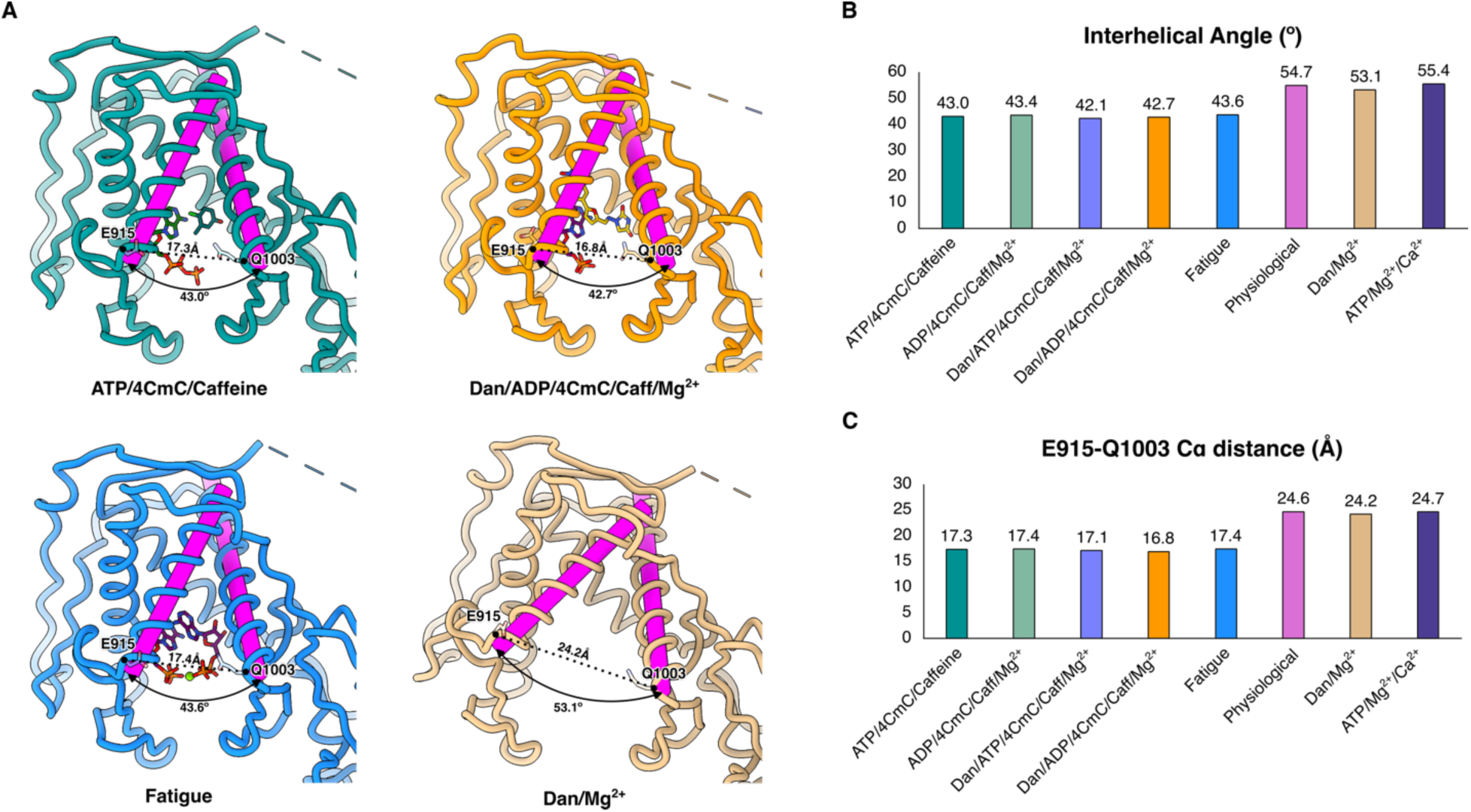
Changes in interhelical angles and Cα distance between E915 and Q1003 in RY12 upon conformational changes. **A**. The magenta cylinder objects were generated in *ChimeraX* for the two α-helices (E915-A934 and P979-A1002) forming the RY12 clamp. The interhelical angles were calculated by measuring the rotational angles between the two cylinders for each dataset. The outermost subdomain of RY12 closes as the ligands bind in the RY12 binding cleft. **B**. Quantification of the interhelical angle measurements. There is ∼11.4° change of the interhelical angles between the apo and the holo states. **C**. Quantification of the E915-Q1003 Cα distance measurements. There is ∼7.3 Å difference between the apo and the holo states.

**Figure S6:**
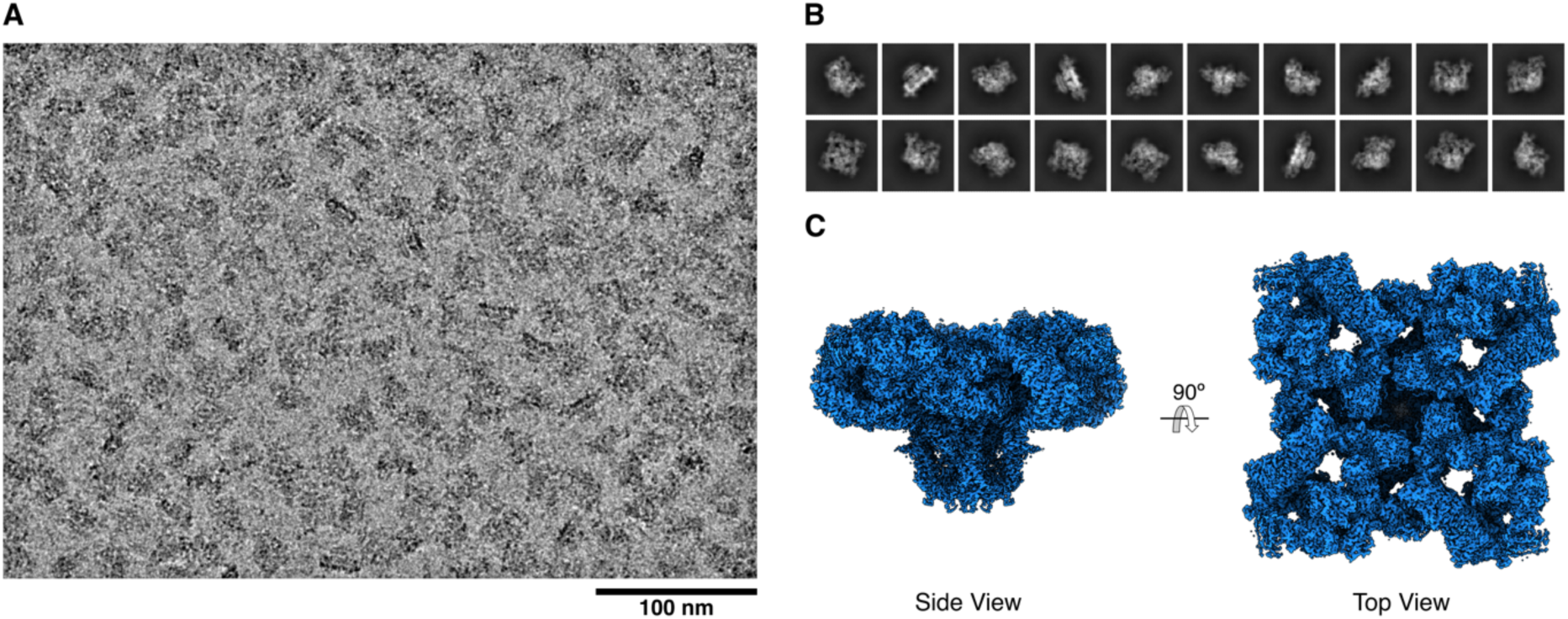
(**A**) A representative cryoEM micrograph of the Fatigue dataset (5,405 micrographs), (**B**) Representative 2D class averages, and (**C**) a combined 3D map was prepared by combining density-modified local maps (Fatigue).

**Figure S7:**
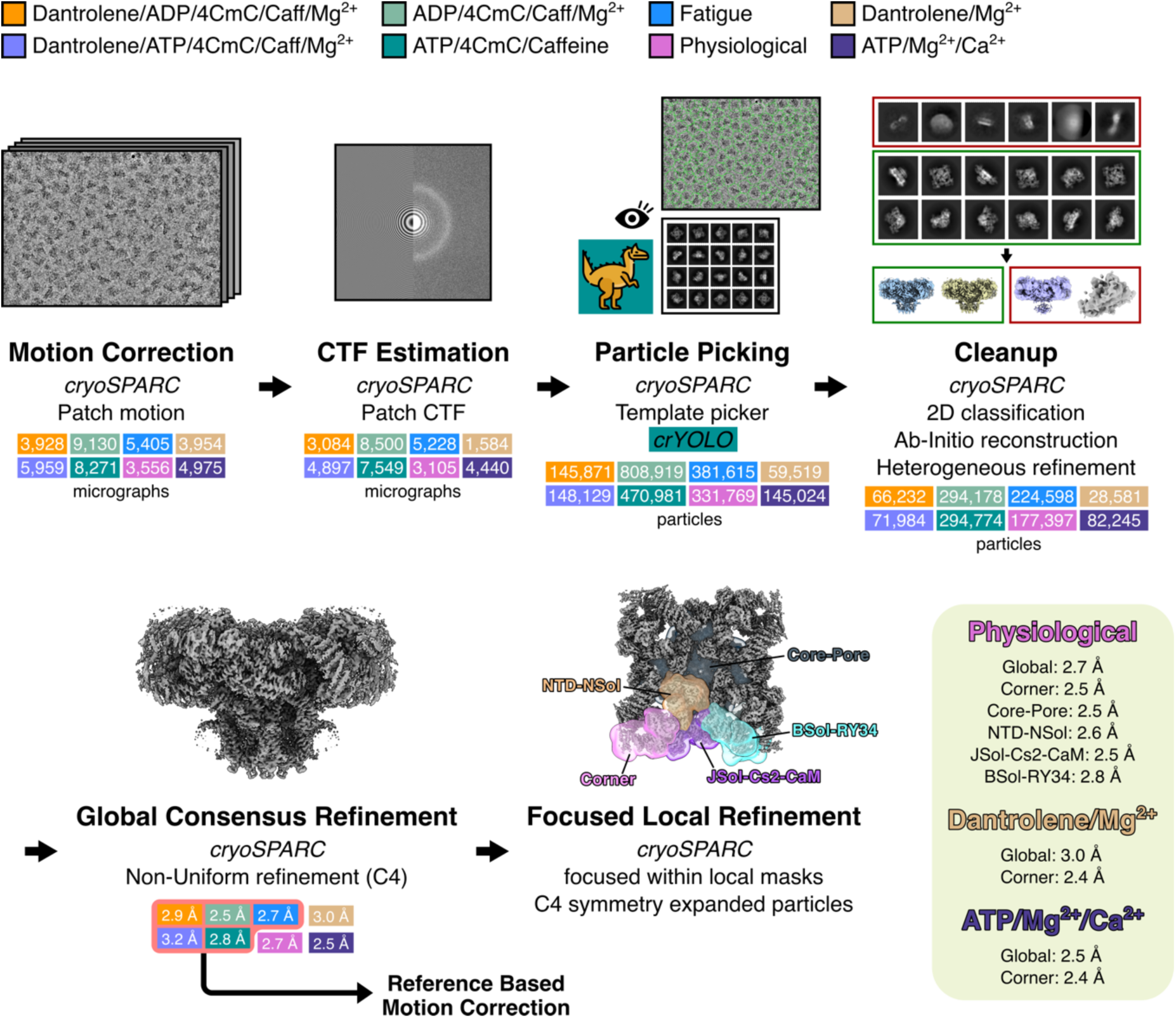
The general cryoEM processing workflow. A flow chart of the generalized cryoEM image processing for the eight color-coded datasets. Particles were picked using *crYOLO* for the ATP/4CmC/Caffeine dataset. Out of the eight datasets, Dan/ADP/4CmC/Caff/Mg^2+^, Dan/ATP/4CmC/Caff/Mg^2+^, ADP/4CmC/Caff/Mg^2+^, ATP/4CmC/Caff, & Fatigue datasets were subjected to reference-based motion correction in *cryoSPARC*.

**Figure S8:**
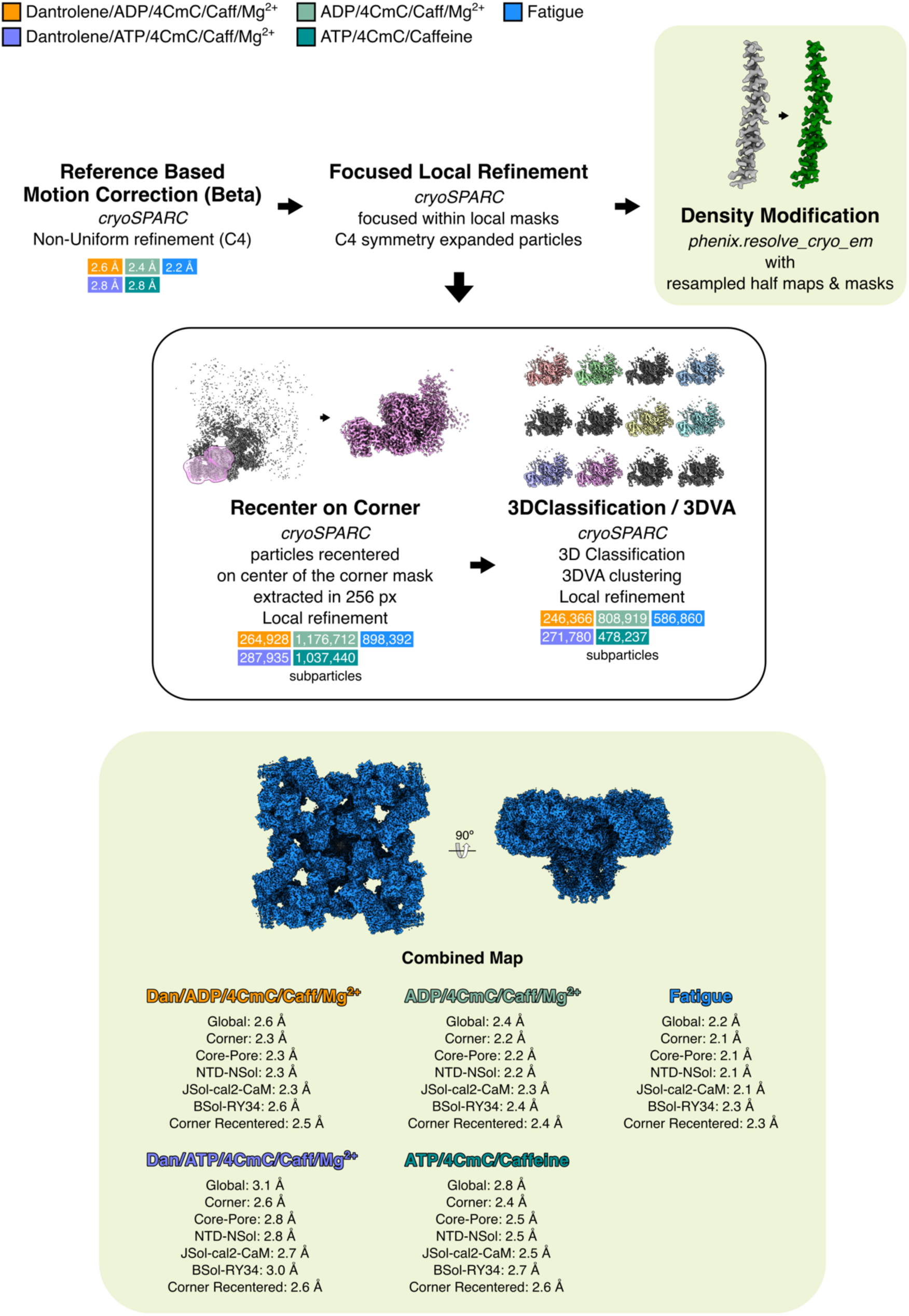
Flowchart of cryoEM image processing for the five datasets (Dan/ADP/4CmC/Caff/Mg^2+^; orange, Dan/ATP/4CmC/Caff/Mg^2+^; orchid, ADP/4CmC/Caff/Mg^2+^; light green, ATP/4CmC/Caff; teal, & Fatigue; blue) that were subjected to Reference-based motion correction, recentering on corner masks and subparticle extraction, 3D Variability Analysis (3DVA) clustering, and 3D classification of the subparticles in *cryoSPARC*. The focused local refinement and density modification steps for the regions other than the corner region were kept the same for all eight datasets.

**Figure S9:**
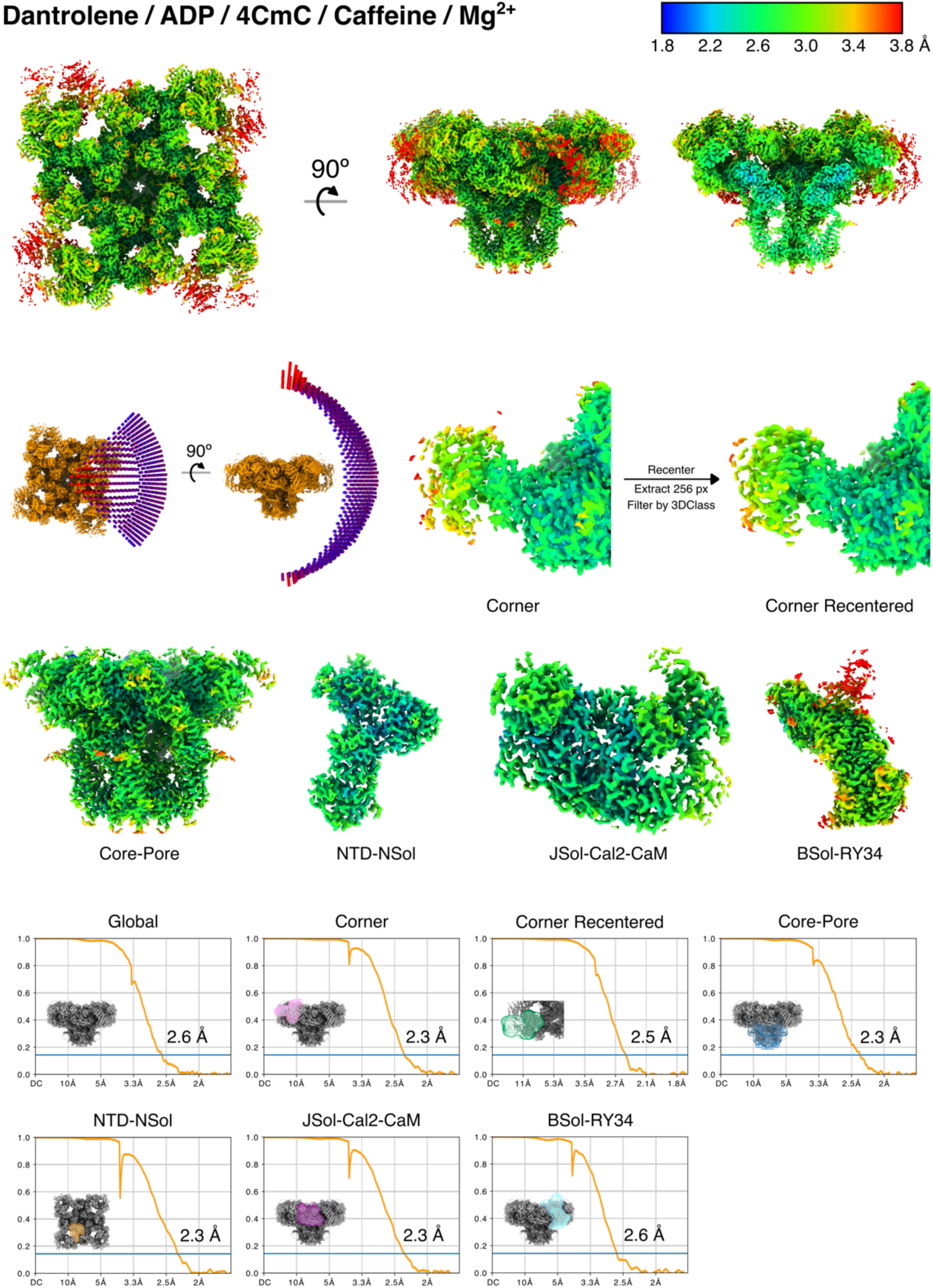
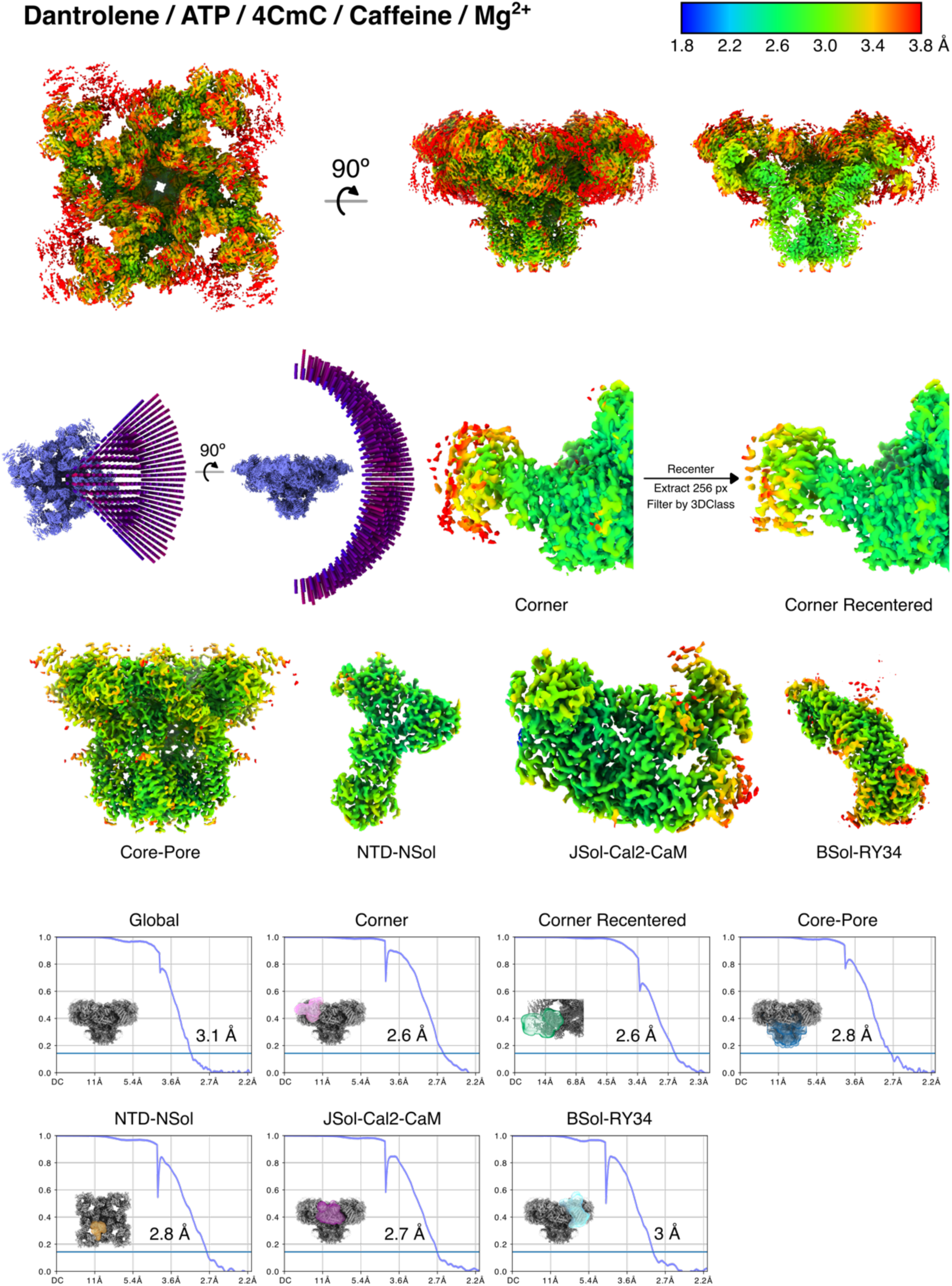
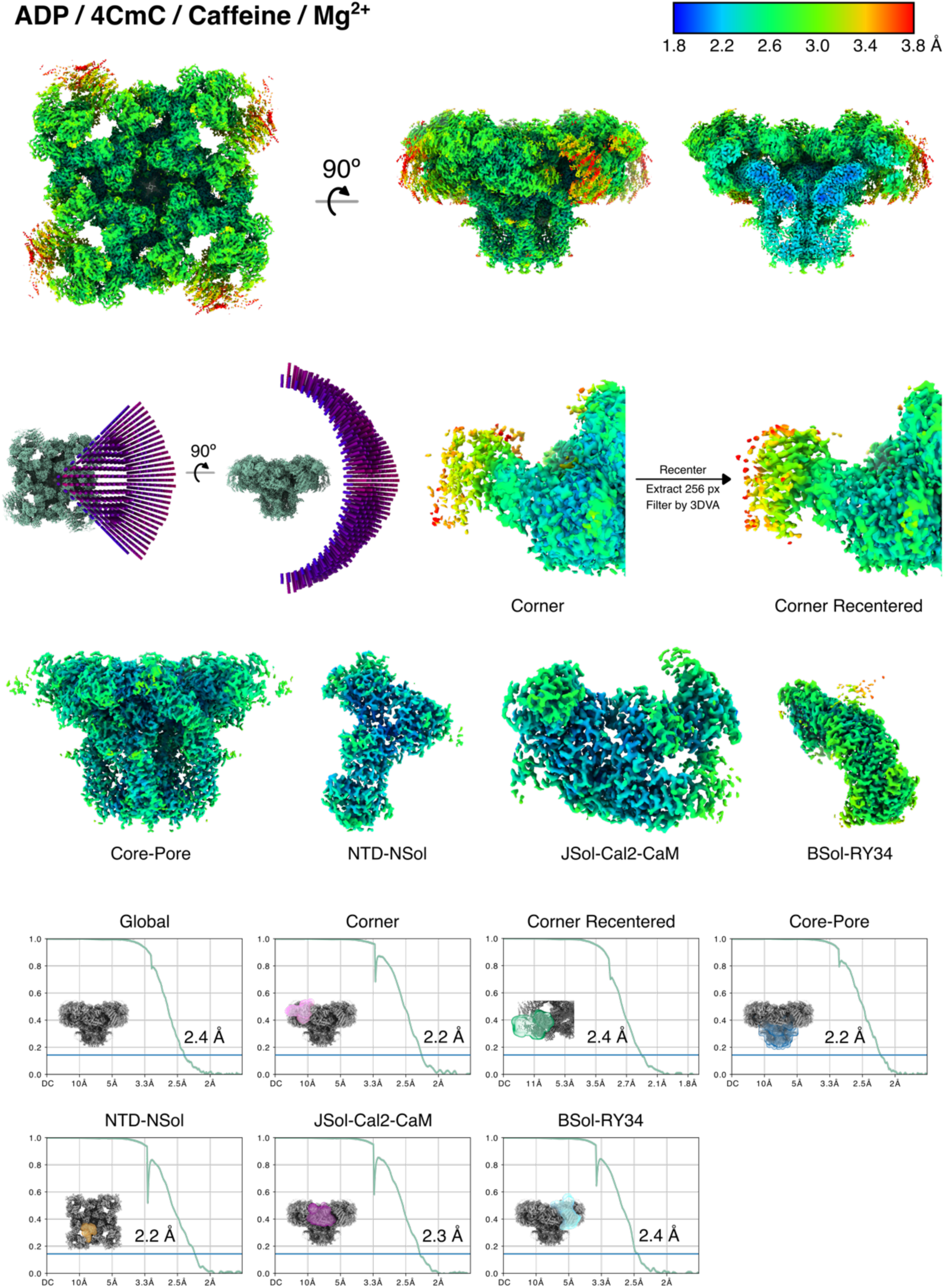
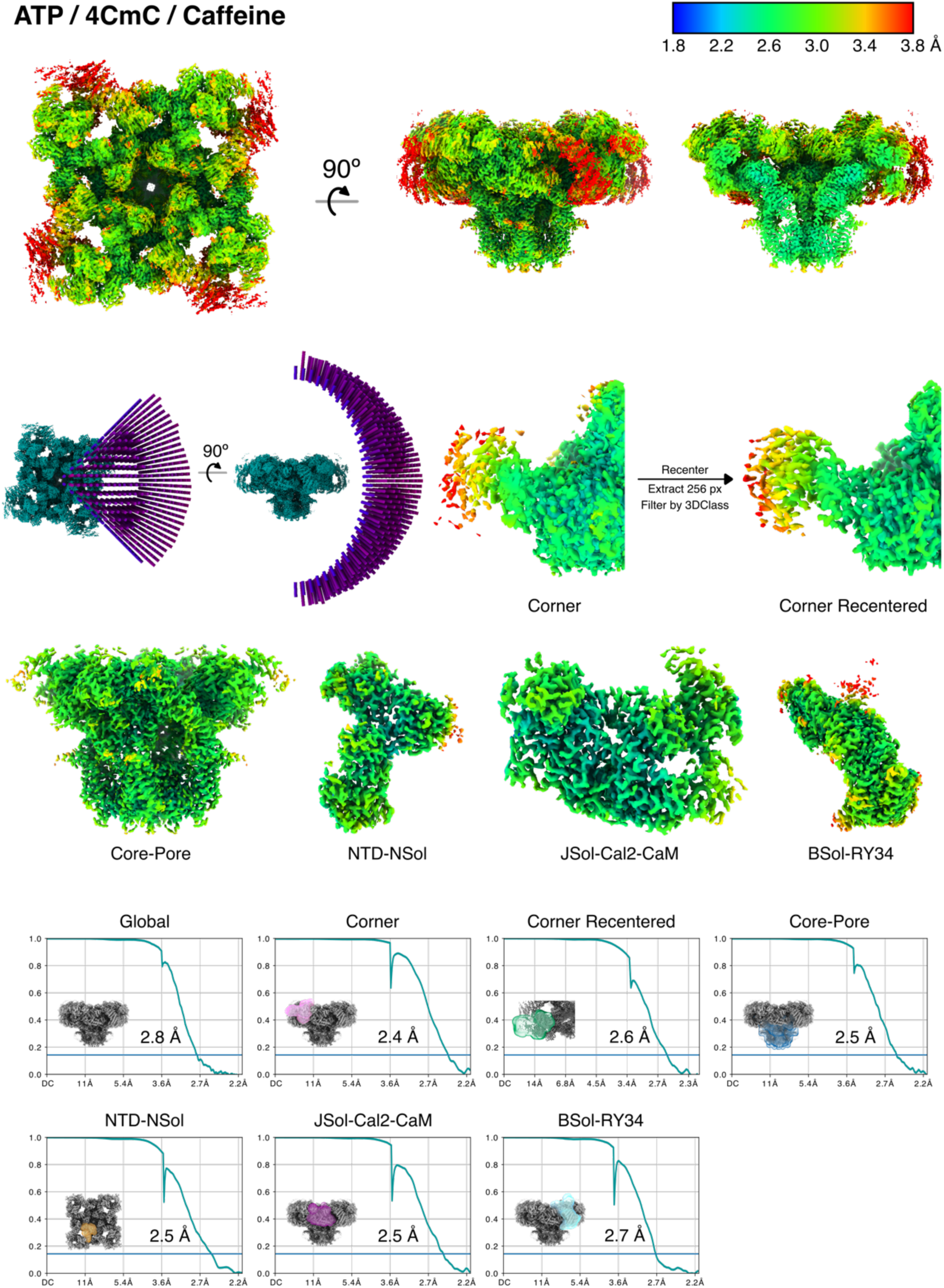
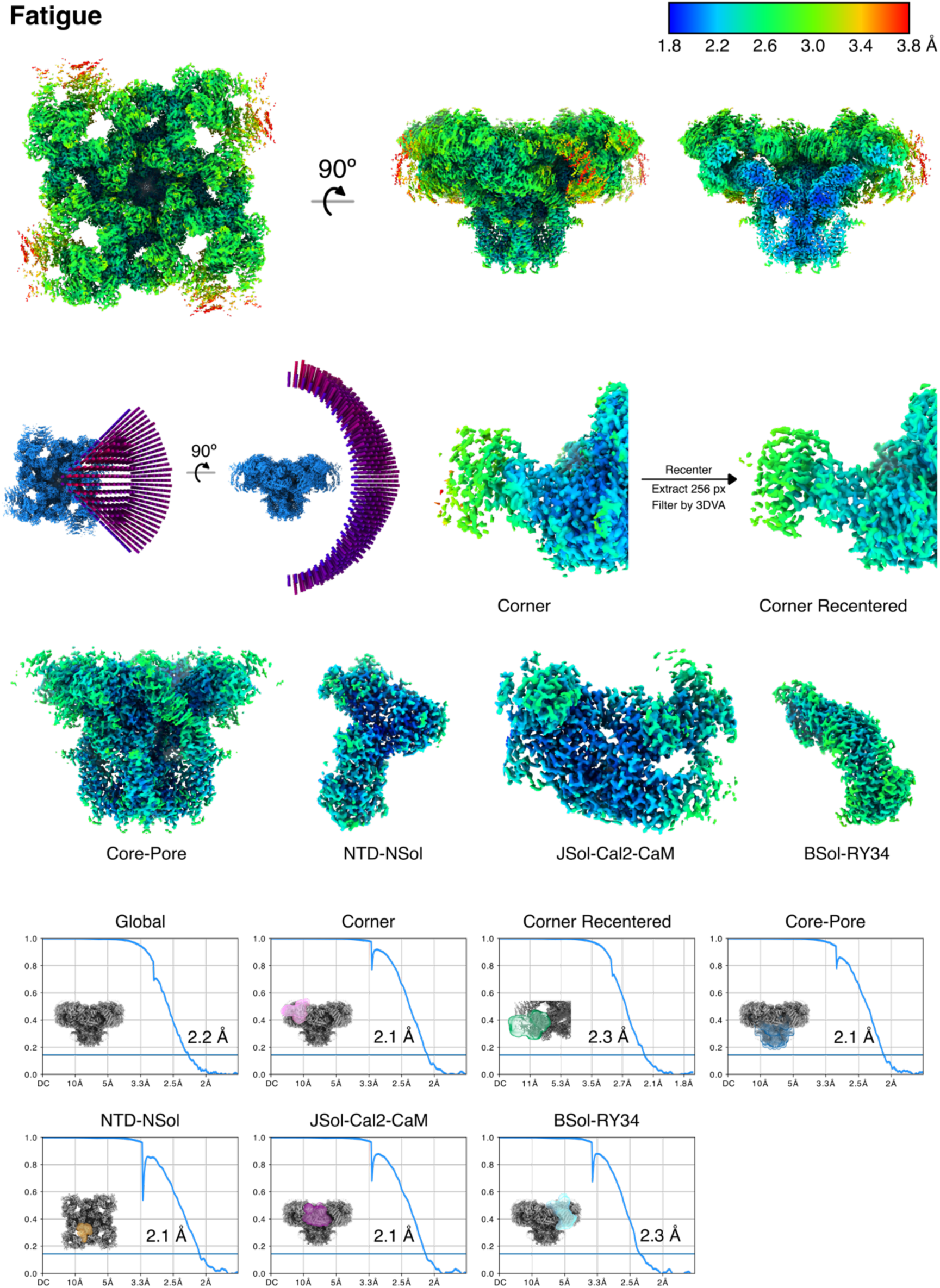
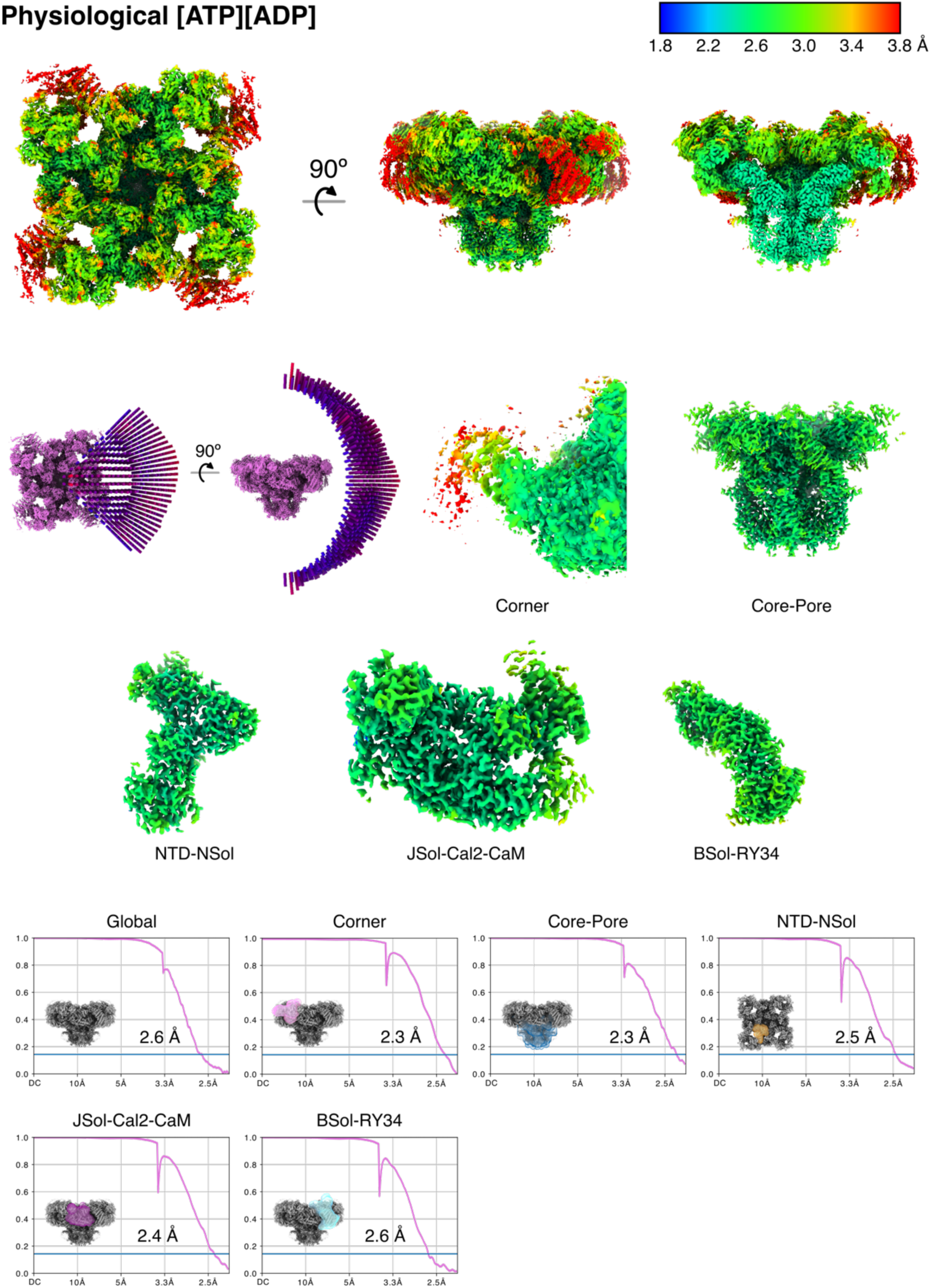
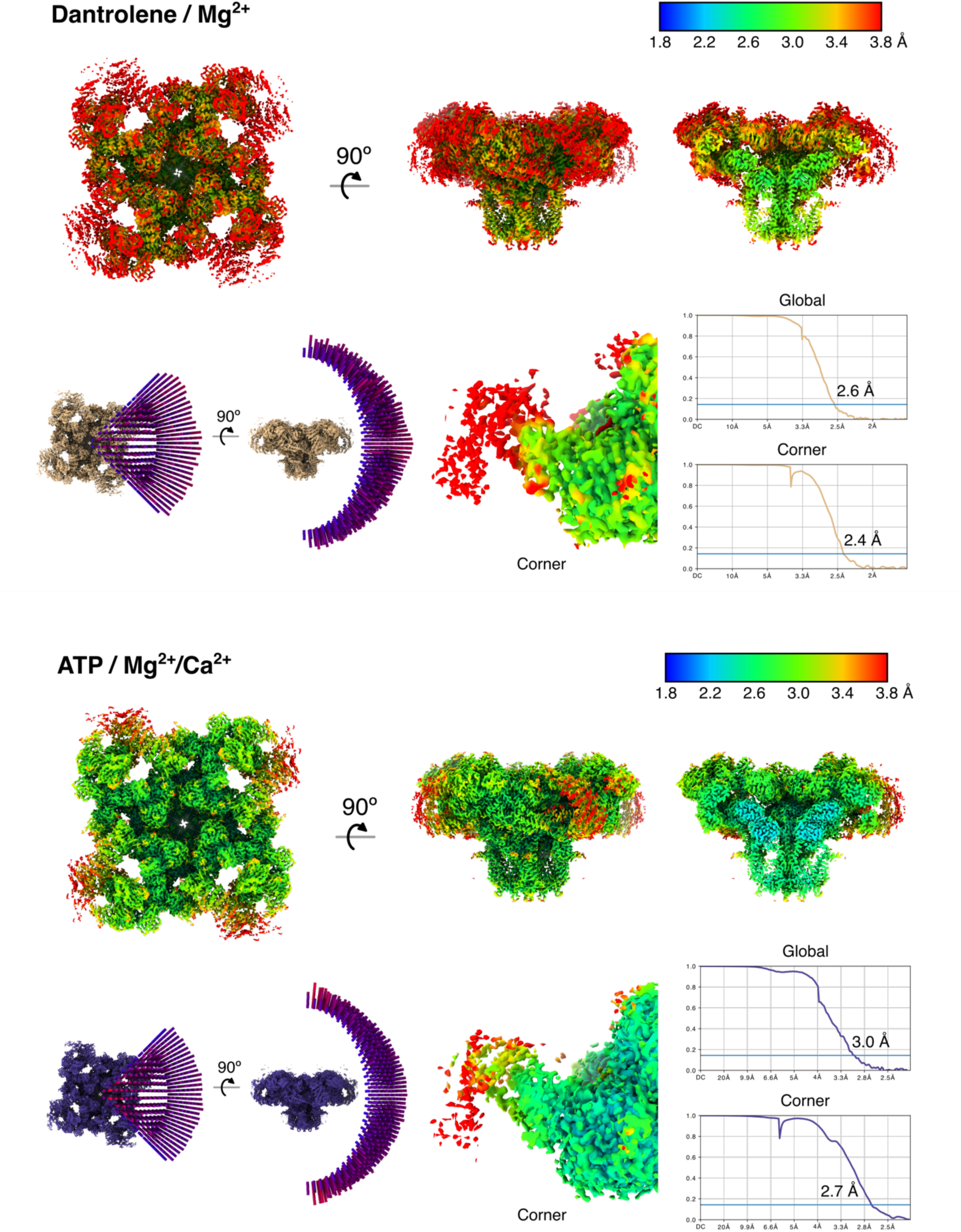
Local resolution and particle orientation plots with FSCs for global and local refinements

**Figure S10:**
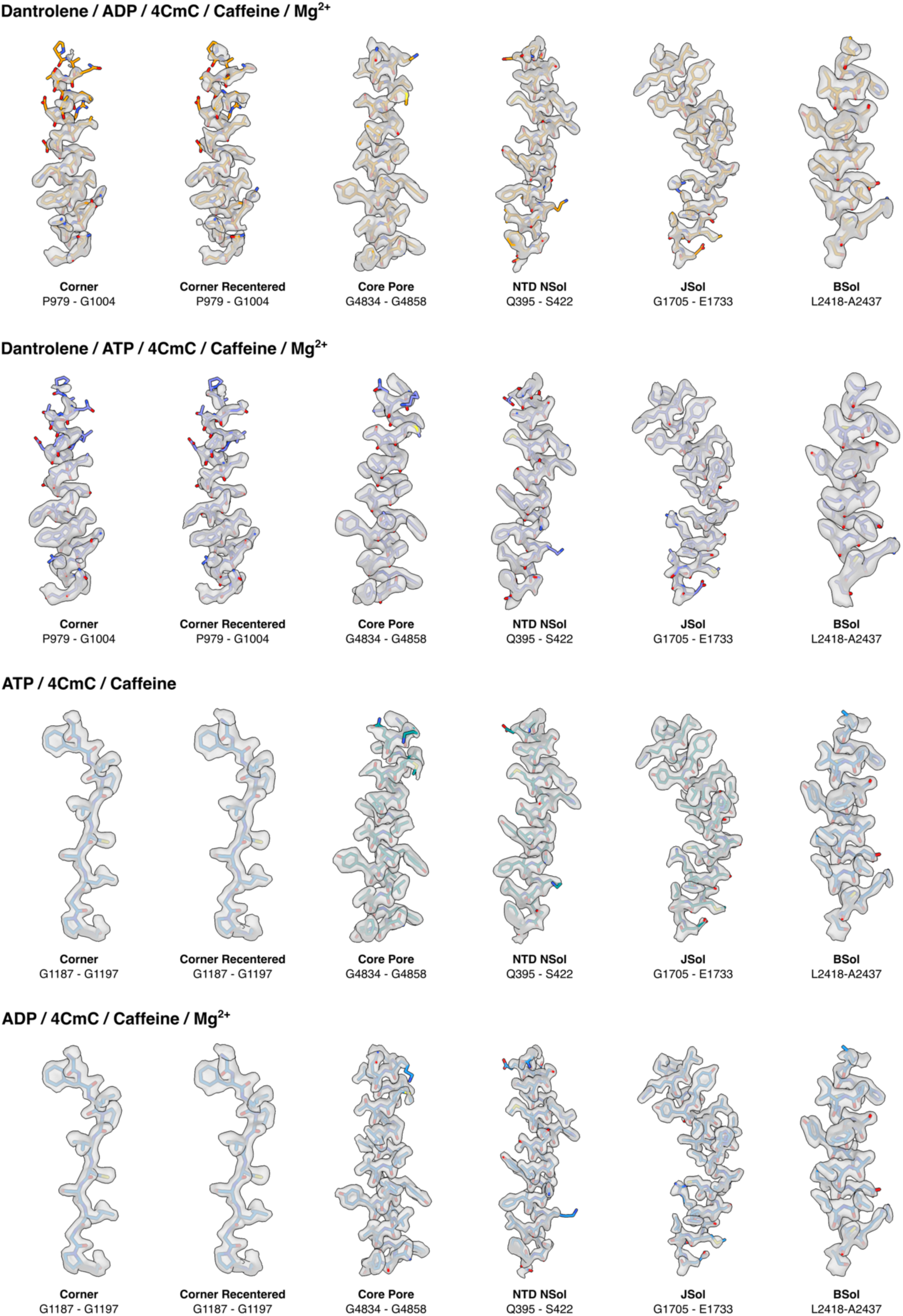
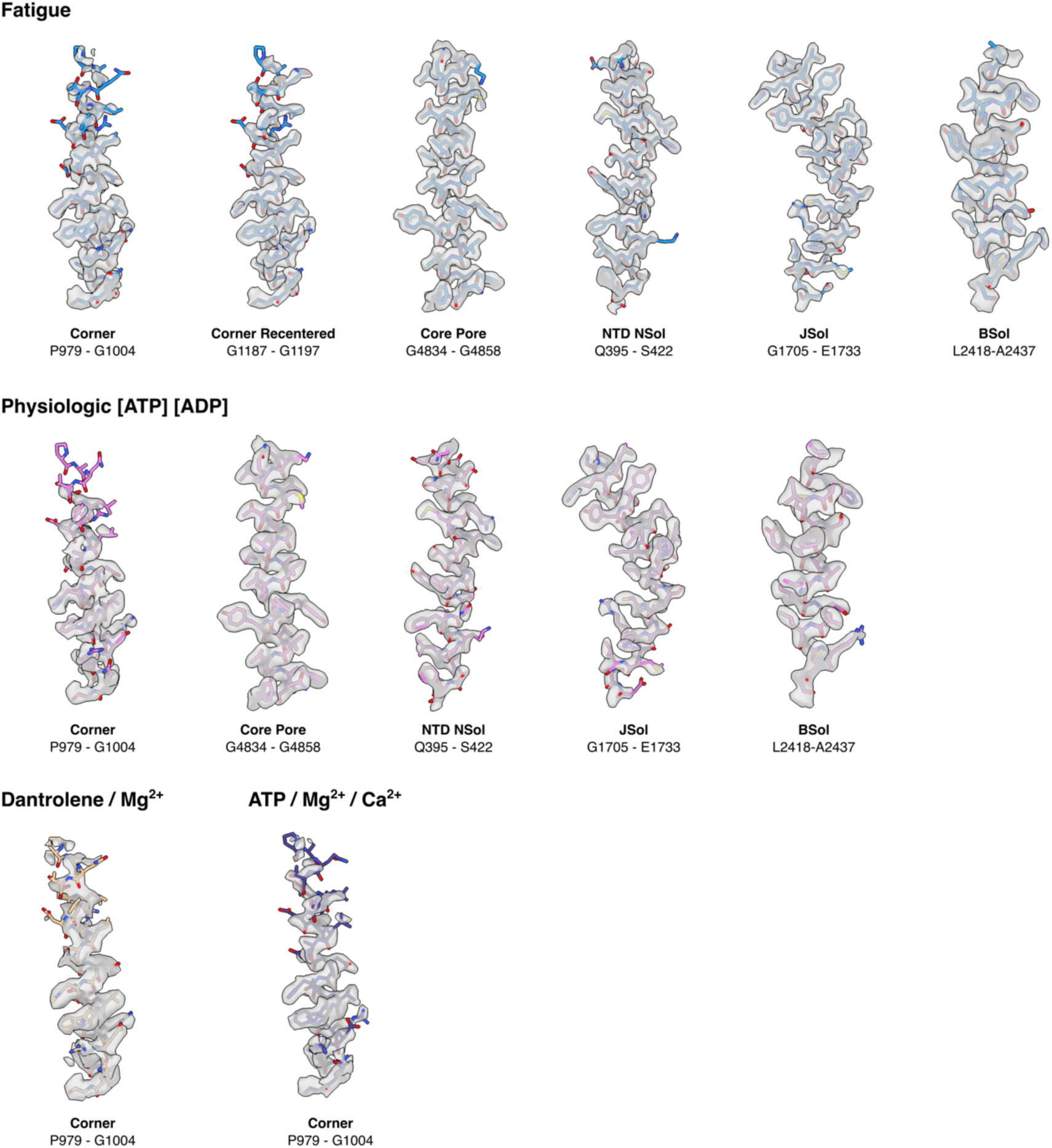
Model/map fit for all local maps of the eight datasets. All maps in the datasets except Dantrolene/Mg^2+^ and ATP/Mg^2+^/Ca^2+^ were subjected to density modification by *resolve_cryo.em* in *Phenix*.

## Methods

### Purification of RyR1 from rabbit skeletal muscle

20-30 g of frozen rabbit skeletal muscle collected from back and thighs of New Zealand White Rabbits (BioIVT) was blended with 150-200 mL of buffer H containing 10 mM Tris Maleate pH 6.8, 1 mM EGTA, 1 mM benzamidine hydrocholoride (Accela ChemBio), and 0.5 mM AEBSF (GoldBio), using a Waring blender for 1.5 minutes. The lysed homogenate was centrifuged at 11,000 x g for 10 minutes to remove large debris, then the supernatant was filtered through three layers of cheesecloth before the second centrifugation at 36,000 x g for 30 minutes. Membrane pellets from the second centrifugation were resuspended in 10 mL buffer S (10 mM HEPES pH 7.4, 0.8 M NaCl, 0.5% CHAPS (GoldBio), 0.1% phosphatidylcholine (Avanti Polar Lipids) solubilized in 10% CHAPS, 1 mM EGTA, 2 mM DTT (GoldBio), 0.5 mM AEBSF, 1 mM benzamidine hydrochloride, and a half of a protease inhibitor tablet (Pierce)) with 10 up and down strokes in a Dounce glass homogenizer. After the 10 strokes, 10 mL buffer D (buffer S without NaCl and protease inhibitors) was added and mixed via another round of 10 up and down strokes. The solubilized membrane homogenate was then subjected to a round of ultracentrifugation at 120,000 x g for 30 minutes. The supernatant was collected and filtered through a 0.2 µm syringe filter before it was loaded onto a 5 mL HiTrap Q HP anion exchange column (Cytiva) pre-equilibrated with buffer W_A_ (10 mM HEPES pH 7.4, 400 mM NaCl, 0.25% CHAPS, 1 mM EGTA, 0.5 mM TCEP (GoldBio), 0.5 mM AEBSF, 1 mM benzamidine, and 0.01% 1,2-dioleoyl-*sn*-glycero-3-phosphocholine (DOPC; Avanti Polar Lipids)). After the filtered supernatant was loaded onto the anion exchange column, contaminant proteins were washed with five column volumes of W_A_. Resin-bound protein was eluted with a linear gradient of 480-520 mM NaCl using buffers W_A_ and W_B_ (buffer W_A_ with 600 mM NaCl). The eluted fractions were checked via SDS-PAGE for bands with high molecular weight corresponding to RyR1, and fractions which contain RyR1 were pooled and subjected to incubation with excess calmodulin and calstabin-2 (Cs2; FKBP12.6) at a 1:40 molar ratio for 1 hour in 4 °C to allow formation of RyR1-Cs2-CaM complexes. Calstabin2 was used to form the complex because it has a higher affinity for RyR1 compared to calstabin1, and it has been shown to sufficiently form a complex with RyR1^8,30^. Calstabin1 (FKBP12) and Cs2 share the same binding site, and both stabilize the closed conformation of RyR1. Concentration of RyR1 in the combined fractions was measured with a Nanodrop One spectrophotometer (Thermo Scientific; 1 abs @ 280 nm = 1 mg/mL). After incubation, the sample is concentrated to 0.5-1 mL using an Amicon centrifugal filter unit with 100 kDa MWCO pore size (Millipore Sigma). The concentrated sample was filtered through a 0.22 µm centrifugal filter unit (Corning) before loading onto a TSKgel BioAssist G4SW_XL_ column (Tosoh Biosciences) pre-equilibrated with buffer E (10 mM HEPES pH 7.5, 0.15 M NaCl, 0.5 mM TCEP, 1 mM EGTA, 0.25% CHAPS, and 0.001% DOPC) for size exclusion chromatography. Fractions were eluted at 0.5 mL/min, and the fractions containing the RyR1-Cs2-CaM complex were pooled and concentrated to 7-9 mg/mL using 100 kDa cut-off Amicon centrifugal filter units. Ligands were added to the concentrated sample before cryoEM grid preparation. ATP, ADP, and free Mg^2+^ concentrations were calculated using *MaxChelator*.^31^ The added ligand concentrations for each condition are summarized in Table 2.

Since dantrolene has poor solubility in H_2_O, dantrolene sodium salt (Sigma) was solubilized in dimethyl fumarate (DMF), then the dantrolene DMF solution was diluted with H_2_O. Briefly, DMF was first purged for 30 minutes with inert nitrogen or argon gas. 12 mM dantrolene in DMF (4.04 mg/mL) was prepared first, then the solution was diluted twenty-fold with H_2_O to prepare a stock solution of 600 *µ*M dantrolene. 100 *µ*M dantrolene was added to the sample to prepare cryoEM grids.

### Cryo-EM grid preparation

UltrAuFoil R0.6/1 Au-300 mesh holey gold grids (Quantifoil) were glow-discharged (PELCO easiGlow), and 3 *µ*L of purified RyR1-Cs2-CaM with ligands was deposited onto the hydrophilized EM grid. The sample was blotted for 4.5 to 5.5 seconds at 4 °C and 100% relative humidity in the chamber of a Vitrobot Mark IV (Thermo Fisher). After blotting, the sample EM grid was plunged into liquid ethane for vitrification. The grids were first screened with either a Glacios electron microscope (Thermo Fisher) equipped with a K3 direct electron detector (Gatan) operated at 200 kV or a Tecnai F20 electron microscope (FEI) equipped with a K2 direct electron detector (Gatan) operated at 200 kV. Screened grids were loaded to a Titan Krios (Thermo Fisher) electron microscope equipped with a K2 (1.06 Å/pixel) or a K3 (0.83 Å/pixel) direct electron detector (Gatan) operated at 300 kV in counting mode using Leginon and Appion data collection software suite.^32,33^ Movies were collected at exposure rates of 8 e^-^/pix/s for 8 seconds in 40 frames (200 ms/frame) (Dan/ATP), 8 e-/pix/s for 10 seconds in 50 frames (200 ms/frame) (ATP/4CmC), or 16 e^-^/pix/s for 2.5 seconds in 50 frames (50 ms per frame) (the rest of the datasets presented) using a defocus range of −0.5 *µ*m to −1.5 *µ*m. A 100 *µ*m objective aperture was used for all collections except the ADP/4CmC dataset. Detailed parameters for all datasets are listed in Table 1.

### Cryo-EM data processing

Most of the image processing steps were run in *cryoSPARC*^21^ unless stated otherwise. Movie frames were aligned with Patch Motion job type in *cryoSPARC*. After CTF estimation with Patch CTF, low quality micrographs with high relative ice thickness, high motion pixel distances, and low CTF fit resolution were removed from each dataset. RyR1 particles were picked with *crYOLO*^34^ or template-based particle picker in *cryoSPARC*. Low-resolution ab-initio models were generated with the initial sets of particles that went through a round of 2D classification. Iterative rounds of 2D classification and heterogeneous refinement further removed particles that do not contribute to high-resolution reconstructions. After cleaning up the particle sets for each dataset, global consensus reconstructions of RyR1 were carried out with Non-Uniform refinement in *cryoSPARC*^35^ with C4 symmetry enforced. For Dan/ATP, Dan/ADP, ATP/4CmC, ADP/4CmC, and fatigue datasets, the particle stacks from Non-Uniform (NU) refinement were subjected to reference-based motion correction (RBMC) in a private beta version (v4.2.2-privatebeta.1) of *cryoSPARC*. RBMC resulted 0.1-0.4 Å nominal resolution improvements for the five datasets. After RBMC, particles were grouped into 25 to 225 groups by similar image shift coordinates recorded during data collection.^36^

In global homogeneous/NU refinement maps, the cytoplasmic domain of RyR1, especially BSol, exhibits a significant degree of motion suggested by smeared or unresolved local map quality. To improve the local resolution, five local soft masks (described in Table 3) were generated using *UCSF ChimeraX*^37^ and *cryoSPARC*. The particle stacks were C4 symmetry-expanded and were subjected to focused refinements in *cryoSPARC* with local masks. These local masks, especially the corner mask (SPRY1- RY12-SPRY2-SPRY3 domains; amino acids 628-1656), were also used for rounds of masked 3D classification without alignment. After an initial local refinement with the corner mask, the C4-expanded particles were re-centered to the centers of the masks at each of the four corner regions of RyR1. These recentered particles were then extracted (or downsized) in a smaller box size (256 px) to shorten processing time. The extracted subparticles were subjected to an initial local refinement then put to 3D variability analysis and 3D classification. Selected particles were then matched back to their original particles before recentering to obtain the maps with the best resolution around the ligands bound in RY12.

The local maps from focused refinements of C4-expanded particles with different local masks around NTD-NSol, Cs2-JSol-CaM, and BSol-RY34 domain regions were aligned to the global consensus maps. The local maps were then combined in *UCSF ChimeraX* by taking the maximum value at each map voxel. The atomic structures of RyR1 were built and fitted in cryoEM maps in *Coot*^38^, and the structures and EM maps were visualized using *UCSF Chimera*^39^ or *ChimeraX*. The local maps were post-processed with density modification, soft masking outside the refinement mask and cropping to a minimal box using *phenix.resolve_cryo_em*^40^ included in the *Phenix* suite.^41^ *LocSpiral*^42^ was used to obtain a map with improved connectivity at the very peripheral region of RY12 for the fatigue dataset.

### Reconstitution of purified RyR1-Cs2-CaM complex into liposomes

We used the on-column liposome formation protocol that was previously described^43^ to incorporate RyR1 into proteoliposomes. Excess calstabin2 and calmodulin were added to RyR1 after the anion-exchange chromatography purification step. Proteins were incubated together overnight in 4 °C. A mixture of lipids (phosphatidylethanolamine (PE):phosphatidylcholine (PC)=5:3, Avanti Polar Lipids) in 10% CHAPS was made and added to the proteins in a 1000:1 lipid:protein ratio. The sample was concentrated to 500 *µ*L with a 100kDa MWCO Amicon centrifugal filter unit and was then loaded onto a hand packed G50 column pre-equilibrated with buffer F (buffer E without CHAPS and DOPC). Proteoliposomal RyR1 was eluted at 0.25 mL/min in buffer F, and fractions corresponding to a single gaussian peak were pooled and checked with SDS-PAGE. The pooled fractions were then concentrated to 5-6 mg/mL, flash frozen with liquid nitrogen, and stored in −80 °C for later usage.

### Single-channel recordings

Purified RyR1 proteoliposomes were fused to planar lipid bilayers formed by painting a lipid mixture of PE and PC in a 5:3 ratio in decane across a 200 *µ*m hole in polysulfonate cups (Warner Instruments) separating 2 chambers. The trans chamber (1.0 mL), representing the intra-SR (luminal) compartment, was connected to the head stage input of a bilayer voltage clamp amplifier. The cis chamber (1.0 mL), representing the cytoplasmic compartment, was held at virtual ground. Asymmetrical solutions used were as follows for the cis solution: 1 mM EGTA, 250/125 mM HEPES/Tris, 50 mM KCl, pH 7.35; and for the trans solution: 53 mM Ca(OH)_2_, 50 mM KCl, 250 mM HEPES, pH 7.35. The concentration of free Ca^2+^ in the cis chamber was calculated as previously described. Purified RyR1 proteins were added to the cis side and fusion with the lipid bilayer was induced by making the cis side hyperosmotic by the addition of 400-500 mM KCl. After the appearance of potassium and chloride channels, the cis side was perfused with the cis solution. At the end of each experiment, 10 *µ*M ryanodine was added to block the RyR channel. Single-channel currents were recorded at 0 mV using a Bilayer Clamp BC-525D (Warner Instruments), filtered at 1 kHz using a Low-Pass Bessel Filter 8 Pole (Warner Instruments), and digitized at 4 kHz. All experiments were performed at room temperature (23°C). Data acquisition was performed by using Digidata 1322A and Axoscope 10.1 software (Axon Instruments). The recordings were analyzed using Clampfit 10.1 (Molecular Devices) and Graphpad Prism software.

### Expression of RyR1-W882A in HEK293T cells

A construct expressing RyR1-W882A was generated by introducing the mutation into fragments of rabbit RYR1, as described previously^9^. HEK293T cells grown in 100- mm dishes with Dulbecco’s Modified Eagle Medium (DMEM) supplemented with 10% (v/v) fetal bovine serum (Invitrogen), penicillin (100 U/ml), streptomycin (100 μg /ml), were transfected with 15 μg per dish of RyR1-W882A cDNA using PEI MAX (Polysciences). Cells were collected 48 hours after transfection. ER vesicles from HEK293 cells expressing RyR1-W882A were prepared by homogenizing cell pellets on ice using a Teflon glass homogenizer with two volumes of solution containing 20 mM tris-maleate (pH 7.4), 1 mM EDTA, 1 mM DTT, and protease inhibitors (Roche). Homogenate was then centrifuged at 8,000xg for 20 min at 4°C. The resulting supernatant was centrifuged at 40,000xg for 40 min at 4°C. The final pellet, containing the ER fractions, was resuspended and aliquoted in 250 mM sucrose, 10 mM Mops (pH 7.4), 1 mM EDTA, 1 mM DTT, and protease inhibitors. Samples were frozen in liquid nitrogen and stored at −80°C.

## Acknowledgements

This work was supported by grants from the National Institutes of Health (R01AR077720 to O.B.C, R01HL142903, R01HL145473, P01HL164319, and R01HL140934 to A.R.M.). A.R.M. is a consultant to, and owns shares in, ARMGO Pharma, Inc., a biotech company targeting RyR channels for therapeutic purposes. The remaining authors declare no competing interests. K.K. would like to thank Francesca Vallese for support and advice. Cryo-EM data were collected at the Columbia University Cryo-Electron Microscopy Center and the Simons Electron Microscopy Center, with technical support from Robert A. Grassucci and Zhening Zhang.

## Author Contributions

Conceptualization, O.B.C., A.R.M, W.A.H, J.F., A.d.G, R.Z. and K.K. Investigation, K.K., H.L., Q.Y., Z.M., and O.B.C. Writing, K.K. and O.B.C.

## Notes

https://www.dropbox.com/scl/fi/elb0vm1evd0y47lnt6mwz/RyR1_maps_and_models.zip?rlkey=0zguve9jievklj1fb9s3mburg&dl=0

